# Genetic variation in haemoglobin alters breathing in high-altitude deer mice

**DOI:** 10.1101/2021.02.23.432531

**Authors:** Catherine M. Ivy, Oliver H. Wearing, Chandrasekhar Natarajan, Rena M. Schweizer, Natalia Gutiérrez-Pinto, Jonathan P. Velotta, Shane C. Campbell-Staton, Elin E. Petersen, Angela Fago, Zachary A. Cheviron, Jay F. Storz, Graham R. Scott

**Affiliations:** Department of Biology, McMaster University, Hamilton, ON, Canada, L8S 4K1; School of Biological Sciences, University of Nebraska, Lincoln, NE, USA, 68588; Divison of Biological Sciences, University of Montana, Missoula, MT, USA, 59812; Department of Ecology and Evolutionary Biology, University of California, Los Angeles, CA, USA, 90095; Department of Biology, Aarhus University, Aarhus, Denmark C, 8000

**Keywords:** evolutionary physiology, hypoxic ventilatory response, hypoxia adaptation, high-altitude adaptation

## Abstract

Physiological systems often have emergent properties but the effects of genetic variation on physiology are often unknown, which presents a major challenge to understanding the mechanisms of phenotypic evolution. We investigated the *in vivo* effects on respiratory physiology of genetic variants in haemoglobin (Hb) that contribute to hypoxia adaptation in high-altitude deer mice (*Peromyscus maniculatus*). We created F_2_ inter-population hybrids of highland and lowland deer mice to test the phenotypic effects of α- and β-globin variants on a mixed genetic background. High-altitude genotypes were associated with breathing phenotypes that enhance O_2_ uptake in hypoxia, including a deeper more effective breathing pattern and an augmented hypoxic ventilatory response. These effects could not be explained by erythrocyte Hb-O_2_ affinity or globin gene expression in the brainstem. Therefore, adaptive variation in haemoglobin can have unexpected effects on physiology that are distinct from the canonical function of this protein in circulatory O_2_ transport.

## INTRODUCTION

High-altitude natives are an exceptional model for understanding the genetic and physiological bases of evolutionary adaptation. Species that are broadly-distributed across altitudes can provide powerful insight into the genetic basis of high-altitude adaptation, because it is possible to examine segregating variation for phenotypes that may contribute to hypoxia tolerance. Recent research has identified many genes that appear to have experienced selection in high-altitude taxa, including genes thought to be involved in O_2_ transport, energy metabolism, and hypoxia signalling (Simonson, 2015; Simonson et al., 2012; Storz and Cheviron, 2021).

However, in most cases, the specific effects of these genetic variants on physiological function are poorly understood. Identifying these functional effects has the potential to uncover novel and adaptive physiological mechanisms, given the growing appreciation that protein variants can have auxiliary effects that are unrelated to the ‘canonical’ function of the protein in question (Marden, 2013a).

Evolved changes in haemoglobin have contributed to hypoxia adaptation in many high-altitude taxa (Storz, 2016). Haemoglobin (Hb) is a tetramer containing two α- and two β-chain subunits, and its O_2_-binding affinity is an important determinant of O_2_ exchange at the lungs and peripheral tissues. Evolved increases in Hb-O_2_ affinity have arisen in many high-altitude taxa, and are typically attributable to amino acid replacements in the α- and/or β-chain subunits that increase intrinsic O_2_-affinity and/or reduce responsiveness to negative allosteric cofactors (e.g. 2,3-DPG in mammals) (Galen et al., 2015; Jendroszek et al., 2018; Natarajan et al., 2018, 2016, 2015b; Projecto-Garcia et al., 2013; Signore et al., 2019; Storz et al., 2010; Tufts et al., 2015; Zhu et al., 2018). An increased Hb-O_2_ affinity is generally assumed to safeguard arterial O_2_ saturation in hypoxia, but this has rarely been tested independent of the other phenotypic differences that covary with Hb-O_2_ affinity in high-altitude taxa. Furthermore, it remains possible that modifications of Hb function contribute to hypoxia tolerance via other physiological mechanisms that are not directly related to circulatory O_2_ transport.

Haemoglobin is not generally thought to be directly involved in the regulation of breathing, but some evidence suggests that tetrameric Hb or its constituent α/β globin monomers may play a role. On the one hand, pharmacological manipulations that increase Hb-O_2_ binding affinity do not generally appear to exert much influence on ventilatory responses to hypoxia (Birchard and Tenney, 1986; Rivera-Ch et al., 1994), nor do some naturally occurring mutant haemoglobins that have altered O_2_ affinity (e.g., Andrew-Minneapolis mutation in the β-globin subunit of humans) (Hebbel et al., 1977). By contrast, affinity-altering mutations in Hbs of mice and sheep are associated with changes in ventilatory sensitivity to O_2_ and/or CO_2_ (Dawson and Evans, 1966; Izumizaki et al., 2003; Shirasawa et al., 2003). Furthermore, deoxygenated Hb can produce *S*-nitrosothiols that contribute to signalling the ventilatory response to hypoxia in rats (Lipton et al., 2001). These findings together suggest that Hb or its globin monomers may play an underappreciated role in the control of breathing, by a mechanism that is not directly associated with the role of Hb in circulatory O_2_ transport. Hb has recently been shown to be expressed in various non-erythroid cells, including neurons and vascular endothelium (Biagioli et al., 2009; Newton et al., 2006; Richter et al., 2009; Schelshorn et al., 2009; Straub et al., 2012), so it is possible that globins expressed in non-erythroid tissues could regulate ventilatory phenotypes.

High-altitude deer mice have evolved an increased Hb-O_2_ affinity relative to lowland conspecifics (Snyder et al., 1982; Storz et al., 2010), and this modification of protein function is associated with increased arterial O_2_ saturation in hypoxia (Ivy et al., 2020; Ivy and Scott, 2017; Tate et al., 2017). Experimental studies have identified the mutations in the α- and β-chain subunits that are responsible for population differences in Hb-O_2_ affinity (Jensen et al., 2016; Natarajan et al., 2015a, 2013; Storz et al., 2010, 2009) and population-genetic analyses on sequence variation in the underlying genes provided evidence that the altitudinal patterning of Hb polymorphism is attributable to a history of spatially varying selection that favors different allelic variants in different elevational zones (Storz and Kelly 2008; Storz et al. 2012). Deer mice have also experienced strong directional selection for increased aerobic capacity for thermogenesis (Hayes and O’Connor, 1999), which has led to evolved increases in maximal rates of O_2_ consumption (V^˙^ O_2max_) in high-altitude populations (Cheviron et al., 2014, 2013; Tate et al., 2020, 2017). In addition, high-altitude mice have evolved an enhanced hypoxic ventilatory response and a deeper breathing pattern (larger tidal volumes but lower breathing frequencies at a given level of total ventilation) under routine conditions, both of which should increase alveolar ventilation and be more effective for gas exchange in hypoxia (Ivy et al., 2020; Ivy and Scott, 2018, 2017).

Here we report an investigation of whether high-altitude Hb variants contribute to evolved changes in control of breathing in high-altitude deer mice. This was achieved using an F_2_ intercross breeding design to isolate the effects of allelic variants of α- and β-globins against an admixed genetic background. We then examined whether the physiological effects of variation in Hb genes resulted from changes in Hb-O_2_ affinity, using efaproxiral (a synthetic drug that acts as a negative allosteric regulator of Hb-O_2_ binding) to pharmacologically reduce Hb-O_2_ affinity *in vivo*. Finally, we used exome sequencing to assess genes surrounding α-globin for linkage equilibrium, and transcriptomics and western blots to evaluate the expression of α- and β-globins in the medulla and pons (brainstem regions responsible for the control of breathing).

## RESULTS

### Effects of globin genotype on physiological responses to acute and chronic hypoxia

When F_2_ hybrids were considered altogether, both acute and chronic hypoxia affected breathing, metabolism, and arterial O_2_ saturation (Table S1, Fig. S1). Adult mice were subjected to acute step-wise decreases in inspired partial pressure of O_2_ (PO_2_) both before and after chronic hypoxia acclimation (6-8 weeks at ∼12 kPa O_2_). Total ventilation increased in response to acute hypoxia due to increases in breathing frequency that offset smaller declines in tidal volume.

**Table 1.**
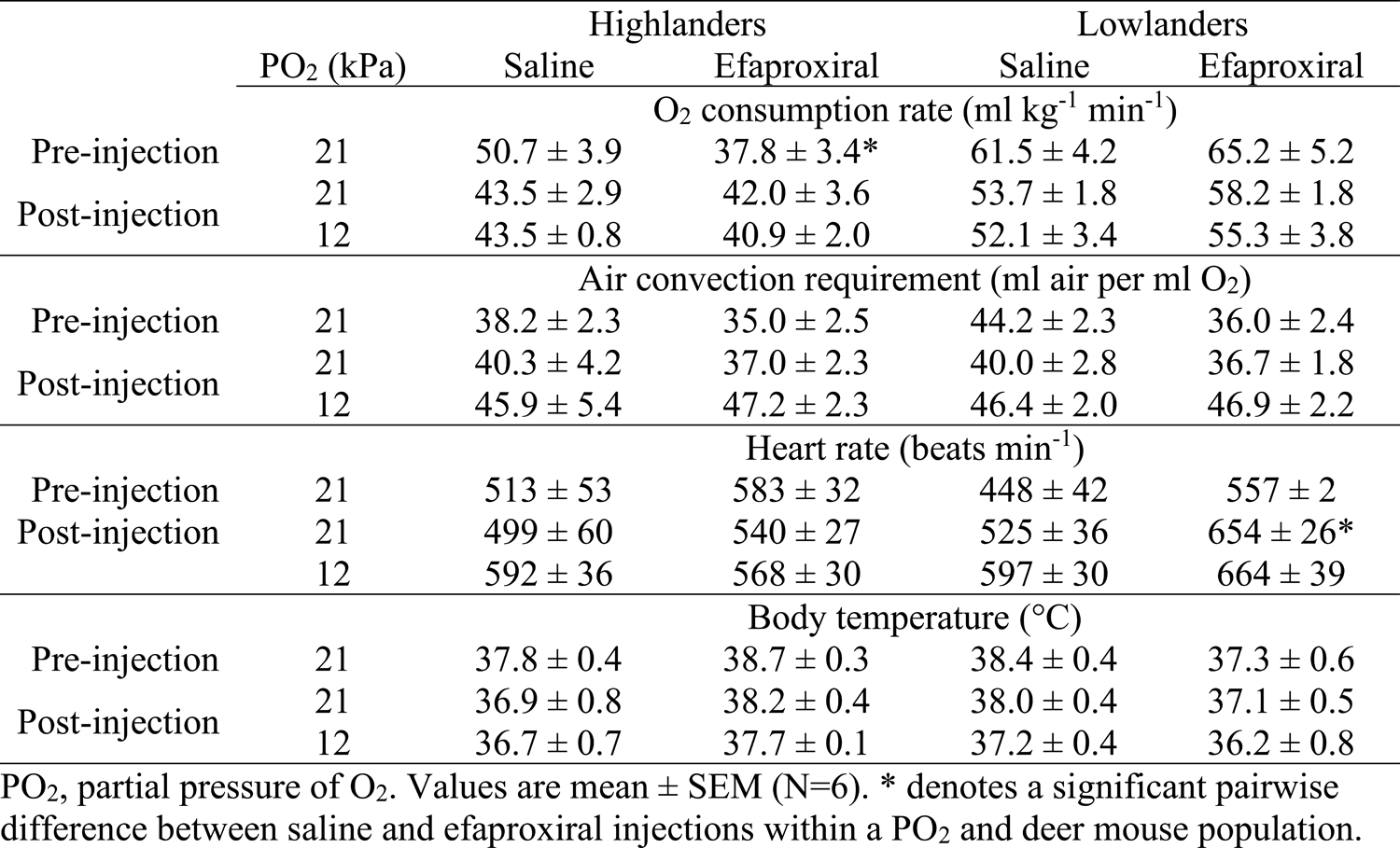
O_2_ consumption rate, air convection requirement, heart rate, and body temperature responses of G_1_ deer mice during manipulation of arterial O_2_ saturation using efaproxiral (200 mg kg^-1^).

Chronic hypoxia augmented these increases in total ventilation, particularly in response to severe acute hypoxia (acclimation environment×inspired PO_2_, P<0.001), which arose largely from significant increases in breathing frequency (environment×PO_2_, P<0.001). Chronic hypoxia also attenuated declines in arterial O_2_ saturation (SaO_2_) (environment effect, P<0.001), O_2_ consumption rate (P=0.009), heart rate (P<0.001), and body temperature (P<0.001) in response to acute stepwise hypoxia compared to normoxia, and increased haematocrit and whole-blood Hb content (Table S2; Fig. S1,S2).

Several of these cardiorespiratory and metabolic phenotypes were associated with α-globin and/or β-globin genotype (Tables S1, S2). Below we discuss the statistically significant effects in the Results, but we include the full suite of measurements for each genotype in Supplementary Figures (Figs. S2,S3,S4). α-globin genotype, but not β-globin genotype, had strong effects on arterial O_2_ saturation (Table S1). SaO_2_ in severe hypoxia was higher in the highland (H) α-globin genotypes compared to the lowland (L) α-globin genotypes in measurements among normoxia-acclimated mice (effect of α-globin genotype, P=0.012; Fig. 1A). However, although chronic hypoxia tended to reduce the decline in SaO_2_ across genotypes (environment effect, P<0.001), this effect was greater in mice with the α^LL^ genotypes, such that the difference in SaO_2_ between genotypes was abolished after hypoxia acclimation (Fig. 1B).

**Figure 1.**
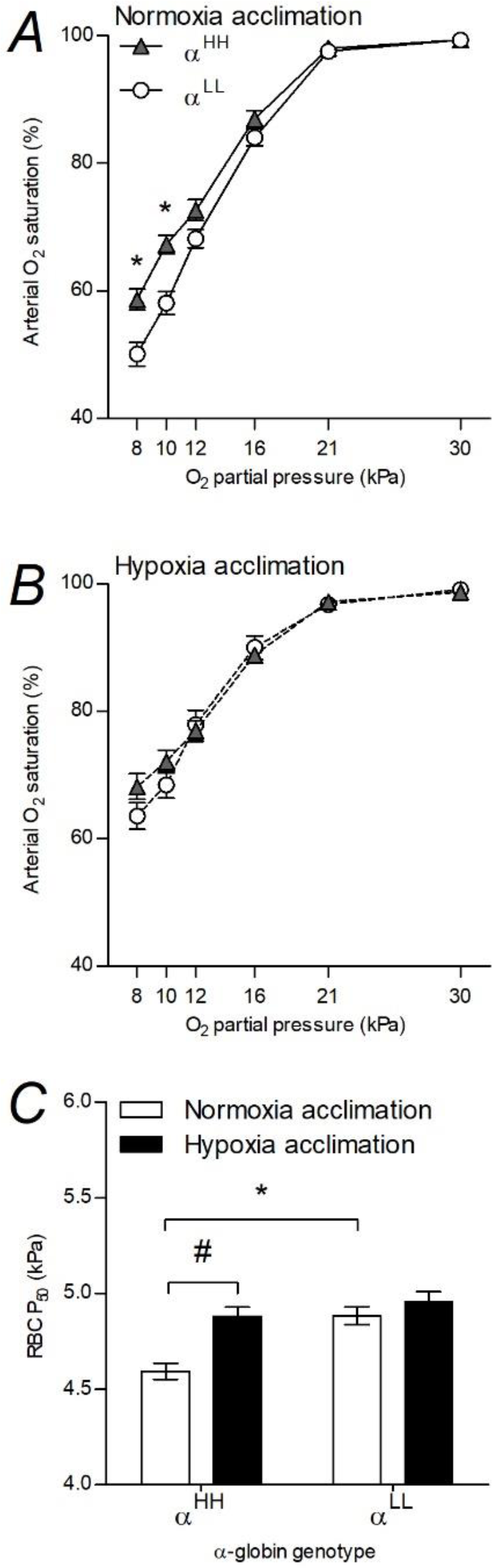
Arterial O_2_ saturation during acute hypoxia (A,B) and red blood cell (RBC) P_50_ (C) were affected by α-globin genotype in F_2_ hybrid deer mice before but not after hypoxia acclimation. Different globin genotypes are shown as superscripts with ‘L’ representing the lowland haplotype and ‘H’ representing the highland haplotype. Values are mean ± SEM (α^HH^, N=17; α^LL^, N=9). * and # denote significant pairwise differences using Holm-Šidák post-tests between α-globin genotypes within an acclimation environment and between acclimation environments within an α-globin genotype, respectively.

This variation in SaO_2_ appeared to be associated with variation in Hb-O_2_ affinity measured in intact red blood cells (RBC), for which there was also a significant effect of α-globin genotype (P<0.001) but not β-globin genotype (Fig. 1C, Table S2). In particular, α^HH^ mice exhibited significantly higher Hb-O_2_ affinity (i.e., lower P_50_) than α^LL^ mice before exposure to chronic hypoxia. However, Hb-O_2_ affinity decreased in response to chronic hypoxia in mice with the α^HH^ genotype, in contrast to mice with the α^LL^ genotype (α-genotype×environment, P=0.035), such that α-globin genotypes were similar after hypoxia acclimation. For measurements of NO metabolites (i.e. nitrite, S-nitrosothiols, iron-nitrosyl and N-nitrosamine derivatives) made after hypoxia acclimation, α-globin genotype affected plasma nitrite concentration (α^HH^ genotype, 0.742 ± 0.070 μM; α^LL^ genotype, 1.289 ± 0.255 μM; P=0.027) but had no effect on plasma concentrations of other NO metabolites (Table S3).

There was also a strong effect of α-globin genotype on breathing pattern, both before and after exposure to chronic hypoxia (Fig. 2, Table S1). In measurements both before and after hypoxia acclimation, mice with the α^HH^ genotype breathed using significantly deeper breaths (α-globin effect, P<0.001) but at a slower frequency (α-globin effect, P<0.001) than mice with the α^LL^ genotype, with no significant differences between α-globin genotypes in total ventilation.

**Figure 2.**
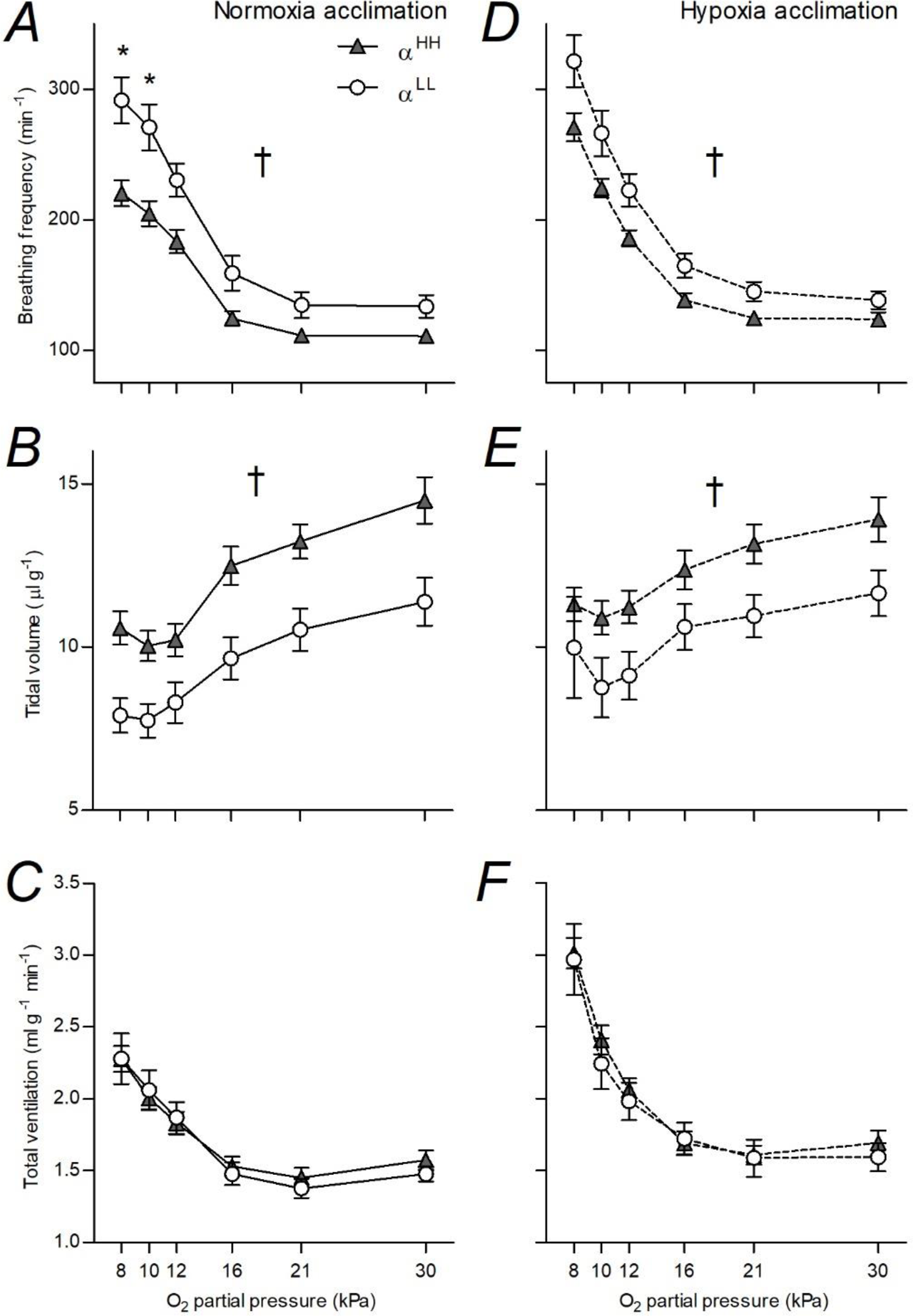
Breathing pattern was altered by α-globin genotype in F_2_ hybrid deer mice both before (A,B,C) and after (D,E,F) hypoxia acclimation. Genotypes and N as in Fig. 1. Values are mean ± SEM. † represents a significant main effect of α-globin genotype, * represents a significant pairwise differences between genotypes within a PO_2_ using Holm-Šidák post-tests.

These differences in breathing pattern persisted across a range of inspired O_2_ levels, from hyperoxia (when Hb was fully saturated with O_2_) to severe hypoxia. Although α-globin genotype affected plasma nitrite levels (Table S3), there was no evidence that differences in NO signalling caused the effects of α-globin genotype on breathing pattern, because manipulation of NO signalling with NO synthase inhibitors had no effect on breathing or metabolism (Fig. S5, Table S4).

β-globin genotype affected the hypoxic ventilatory response, as reflected by a significant interaction between β-globin genotype and inspired PO_2_ on total ventilation (P=0.009). β-globin genotype had a particularly strong influence in normoxia-acclimated mice that were homozygous for highland α-globin (Fig. 3). Among these normoxia-acclimated mice, those that were homozygous for highland β-globin had higher total ventilation than both heterozygotes and lowland homozygotes at 12 kPa O_2_, and higher total ventilation than heterozygotes in more severe levels of acute hypoxia (Fig. 3A). However, these differences between β-globin genotypes disappeared after hypoxia acclimation (Fig. 3B), potentially because the effects of hypoxia acclimation were greater in heterozygotes and lowland homozygotes. In contrast, β-globin genotype had no significant effects on O_2_ consumption rate either before or after hypoxia acclimation (Fig. 3C,D; Table S1).

**Figure 3.**
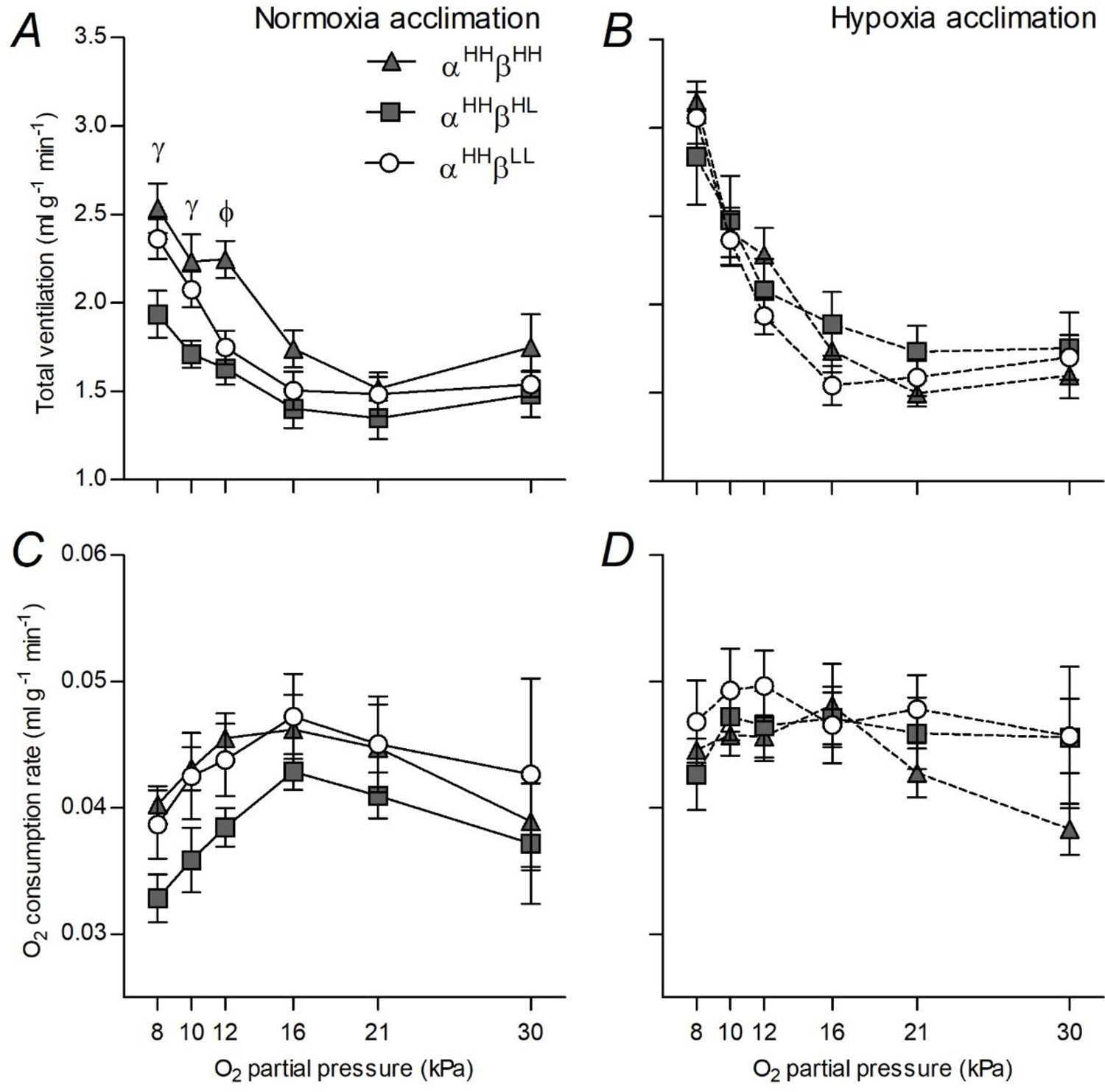
β-globin genotype affected total ventilation in F_2_ hybrid deer mice before (A) but not after (B) hypoxia acclimation, without any significant effects on O_2_ consumption rate (C,D). Genotypes defined in Fig. 1. Values are mean ± SEM (α^HH^β^HH^, N=5; α^HH^β^HL^, N=5; α^HH^β^LL^, N=7). γ and ϕ represent significant pairwise differences within a PO_2_ between α^HH^β^HH^ and α^HH^β^HL^, or between α^HH^β^HH^ and both α^HH^β^HL^ and α^HH^β^LL^, respectively (Holm-Šidák post-tests).

Body temperature was affected by both α-globin (α-globin effect, P=0.007) and β-globin (β-globin effect, P=0.037) genotypes (Table S1; Fig. S4). T_b_ in normoxia was similar across genotypes, ∼36-38 °C on average. Both the magnitude of T_b_ depression as well as the PO_2_ at which T_b_ depression occurred varied among genotypes, but the magnitude of T_b_ depression tended be reduced after hypoxia acclimation for all genotypes. However, neither α-globin nor β-globin genotype had significant effects on O_2_ consumption rate before or after hypoxia acclimation (Fig. S4; Table S1).

To examine the possibility that genotypes at loci other than globins might explain the observed phenotypic differences, we determined the extent of linkage disequilibrium around α-globin due to its strong apparent effect on breathing pattern (see Supplementary Materials). Based on analysis of 23 sequenced nuclear exomes, we identified an ∼11.65 Mb region containing 116 genes (including α-globins) that appear to be in the same linkage group (Fig. S6; Tables S5,S6). A subset of 43 genes contained at least one missense SNP (Table S7). Gene ontology (GO) enrichment analysis of all 116 linked genes within the identified region identified a number of significant functional categories, most notably terms related to gamma-aminobutyric acid (GABA) signaling (Table S8). However, when GO enrichment analysis was performed for only the linked genes that contained at least one missense SNP and excluding α-globins, the only significant categories detected were molecular functions related to serine hydrolase activity (GO:0017171, 0008236, 0004252; P_adj_ = 0.018 for all), such that there was no enrichment of any biological processes related to the control of breathing or respiration, lungs, carotid bodies, medulla oblongata, etc. Therefore, although it remains possible that variants other than missense mutations (e.g., in regulatory regions of genes involved in GABA signaling) might contribute to the observed differences in breathing pattern, genetic variation in α-globin remains the most likely cause.

### Effects of manipulating arterial O_2_ saturation on breathing pattern

We next sought to examine whether the effects of globin genotype on respiratory phenotypes stem from variation in Hb-O_2_ affinity. We used efaproxiral – a synthetic drug that acts as a negative allosteric regulator of Hb-O_2_ binding – to reduce Hb-O_2_ affinity of captive-bred deer mice from high- and low-altitude populations *in vivo*. This treatment was expected to manifest as a reduction in arterial O_2_ saturation in acute hypoxia. Indeed, efaproxiral reduced arterial O_2_ saturation in high-altitude deer mice in hypoxia (Fig. 4A) and in low-altitude deer mice in both normoxia and hypoxia (Fig. 4B) compared to saline controls (P<0.001 for treatment effect and treatment×PO_2_ interaction; Table S9). The magnitude of the effect of efaproxiral on arterial O_2_ saturation differed between populations (population×treatment, P=0.002; population×treatment×PO_2_, P=0.048), driven by larger effects in lowlanders than in highlanders. However, efaproxiral had no consistent effects on breathing frequency, tidal volume, or total ventilation in each population (no significant treatment or treatment×PO_2_ effects) (Fig. 4, Table S9). Efaproxiral also had no consistent effects on oxygen consumption rate, air convection requirement, or body temperature (Tables 1, S9). However, efaproxiral did affect heart rate, as reflected by a significant treatment×PO_2_ interaction (P=0.005) that was driven primarily by increased heart rates in lowlanders after efaproxiral injection (population×treatment, P=0.012; Tables 1, S9). Therefore, our treatment was successful in reducing arterial O_2_ saturation and leading to potential compensatory changes in heart rate but it had no effect on the control of breathing, in stark contrast to the differences observed between globin genotypes.

**Figure 4.**
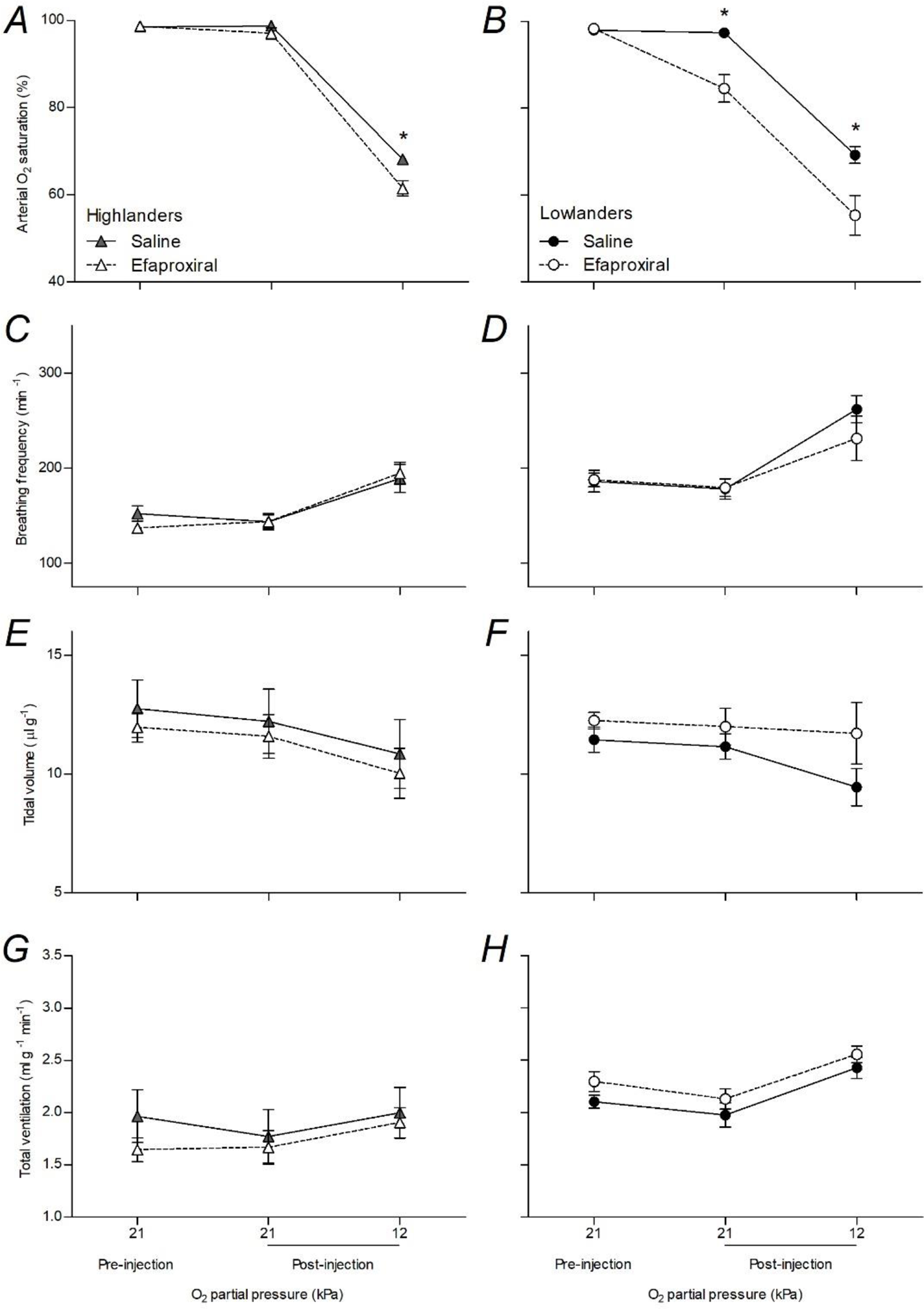
Efaproxiral treatment to reduce haemoglobin-O_2_ affinity reduced arterial O_2_ saturation but did not influence breathing pattern in highland or lowland populations of deer mice. Values are mean ± SEM (N=6). * represents a significant pairwise differences between saline and efaproxiral (200 mg kg^-1^) treatments within a PO_2_ using Holm-Šidák post-tests.

### Globin expression and transcriptomics of the brainstem

We next sought to examine whether the effects of globin genotype on respiratory phenotypes could have arisen as a result of globin expression in key brainstem sites of ventilatory control, and whether these effects were associated with changes in gene expression in the medulla. High-throughput sequencing of medulla transcriptomes from F_2_ hybrids with α^HH^β^HH^ (n=5) and α^LL^β^HL^ genotypes (n=5), sampled in chronic hypoxia, detected 16,082 genes for which there were at least 20 normalized reads per individual on average. Neither α-globin transcripts (*Hba*) nor β-globin transcripts (*Hbb*) were detected in this transcriptomic data set, as these globins were expressed at very low levels that did not pass our filter. Furthermore, neither α-globin nor β-globin peptide chains were detected using Western blots of perfused brainstem tissue from the medulla and pons, supporting that these globins are not expressed at biologically meaningful levels (Fig. S7). Therefore, the effects of globin genotype on breathing were not a result of globin expression in key brainstem sites of respiratory control.

Differential expression analysis of medulla transcriptomes detected no significant differences in gene expression between α^HH^β^HH^ mice and α^LL^β^HL^ mice after transcriptome-wide correction for false discovery rate (FDR). Furthermore, only four genes approached significance – *Il1rap*, *Trpv3*, *Decr2*, and *Cntrob* (P = 0.050 for all) – all of which had reduced expression in α^HH^β^HH^ mice compared to α^LL^β^HL^ mice (Table S10). Weighted gene co-expression network analysis (WGCNA) identified 69 gene expression modules across the medulla transcriptome, but they were not differentially expressed between globin genotypes (Table S11). Therefore, the effects of globin genotype on control of breathing do not appear to be associated with medulla-wide changes in gene expression.

## DISCUSSION

Evolved changes in Hb-O_2_ affinity have contributed to hypoxia adaptation in numerous high-altitude mammals and birds and is often assumed to confer a physiological benefit by safeguarding arterial O_2_ saturation in hypoxia. However, the possibility that adaptive modifications of Hb function might contribute to other physiological processes that are not directly related to circulatory O_2_ transport had been largely unexplored. Here, we show that allelic variants of the α-globin and β-globin genes in high-altitude deer mice lead to changes in control of breathing that augment alveolar ventilation in hypoxia. These effects did not appear to result from changes in arterial O_2_ saturation or Hb-O_2_ affinity, nor were they associated with non-erythroid expression of globins in key brainstem sites of ventilatory control or with appreciable changes in gene expression in the medulla. Nevertheless, our results suggest that allelic variation in Hb genes may affect multiple respiratory phenotypes, and may contribute to environmental adaptation via physiological mechanisms that are not commonly ascribed to this protein.

α-globin genotype had a strong influence on breathing pattern, in which α^HH^ mice breathed deeper but less frequently, a change that likely augments alveolar ventilation. Observed differences in breathing pattern between α-globin genotypes can completely account for previously documented differences that distinguish highland deer mice from lowland conspecifics and a closely related lowland congener, *Peromyscus leucopus*. These differences in breathing pattern are observed in wild mice as well as mice raised for one or two generations in captivity in normoxia (Ivy et al., 2020; Ivy and Scott, 2018, 2017). Such changes in breathing pattern have also evolved in the high-altitude bar-headed goose (Scott et al., 2007), which also possesses an α-globin variant that contributes to an evolved increase in Hb-O_2_ affinity (Natarajan et al., 2018), suggesting that there may be a causal association between α-globin genotype and breathing pattern in other high-altitude taxa.

β-globin genotype appeared to affect the ventilatory response to hypoxia among normoxia acclimated mice, with α^HH^β^HH^ mice exhibiting higher total ventilation than α^HH^β^HL^ and α^HH^β^LL^ mice at 12 kPa O_2_. These findings mirror those in lab-strain mice possessing the Hb Presbyterian β-globin mutation, which is associated with a reduced Hb-O_2_ affinity and an attenuated hypoxic ventilatory response (Izumizaki et al., 2003). Differences in β-globin genotype could thus, in addition to α-globin, contribute to the increases in total ventilation in highland deer mice compared to lowland deer mice and *P. leucopus* that we have observed among mice acclimated to normoxia (Ivy et al., 2020; Ivy and Scott, 2018, 2017). However, these effects of β-globin genotype were abolished after hypoxia acclimation, suggesting that β-globin may influence the effects of chronic hypoxia on control of breathing, a process termed ventilatory acclimatization to hypoxia (VAH). If so, β-globin may contribute to the attenuation of VAH that we have previously observed in highland deer mice (Ivy and Scott, 2018, 2017).

Effects of globin variants on control of breathing did not appear to result from variation in Hb-O_2_ affinity or arterial O_2_ saturation. Treatment of deer mice from highland and lowland populations with efaproxiral to reduce Hb-O_2_ affinity had no effect on total ventilation or breathing pattern in normoxia or hypoxia. There were differences in the magnitude of the effects of efaproxiral on arterial O_2_ saturation and heart rate between populations, possibly because the normally lower Hb-O_2_ affinity of lowlanders (Ivy et al., 2020) made them more susceptible to impairments in pulmonary O_2_ loading upon further reduction in affinity, but efaproxiral had no effect on breathing in either population. Furthermore, the differences in breathing pattern between α^HH^ and α^LL^ mice persisted in hyperoxia (30 kPa O_2_) when blood O_2_ tension was well above that needed to fully saturate Hb with O_2_. These findings suggest that the effects of globin variants on breathing arise from mechanisms that are not directly associated with the role of Hb in circulatory O_2_ transport. These mechanisms do not appear to involve non-erythroid expression of globins in key brainstem sites of ventilatory control, nor are they associated with medulla-wide changes in gene expression, and most likely involve actions of globins at other sites.

The effects of genetic variation in α-globin on arterial O_2_ saturation were contingent upon acclimation environment, consistent with other recent findings (Wearing et al., 2020). Mice that were homozygous for highland α-globin (α^HH^) maintained higher arterial O_2_ saturation in hypoxia and had higher Hb-O_2_ affinity than lowland homozygous mice (α^LL^) when comparisons were made among normoxia-acclimated mice, but these differences were abolished after hypoxia acclimation. This discrepancy could be explained by differences in sensitivity to 2,3-DPG, a key negative allosteric regulator of Hb-O_2_ binding in mammalian erythrocytes that typically increases in concentration after hypoxia acclimation (Lenfant et al., 1968). Indeed, previous studies of O_2_-binding properties of stripped haemoglobin suggest that 2,3-DPG sensitivity is greater in high-altitude populations of deer mice when measured in the presence of physiologically relevant concentrations of Cl^-^ (Storz et al., 2010). Therefore, highland homozygotes could have been more sensitive to the increases in red cell 2,3-DPG concentration that may have occurred with hypoxia acclimation, and have thus exhibited a more pronounced decrease in Hb-O_2_ affinity and a less pronounced increase in arterial O_2_ saturation in hypoxia.

The unexpected association between Hb genotype and control of breathing is especially intriguing in light of population genetic evidence for altitude-related selection on the α- and β-globin genes in deer mice (Storz and Kelly 2008; Storz et al. 2009, 2012). The adaptive relevance of Hb-O_2_ affinity is well-established in high-altitude vertebrates, but the present findings force us to consider the possibility that allelic variation in Hb function may affect a broader diversity of physiological processes than previously assumed. Recent studies suggest that Hb functions not just as an O_2_ carrier, but also as an O_2_ sensor and O_2_-responsive transducer of nitric oxide (NO) vasoactivity in the microcirculation, thereby contributing to hypoxic vasodilation that helps match perfusion to tissue O_2_ demand (Jensen, 2009; Storz, 2018; Zhang et al., 2016, 2015). The results reported here suggest that the physiological effects of Hb may even transcend circulatory O_2_ transport, with direct or indirect effects on control of breathing. The next step is to identify and characterize the causal mechanism underlying the unexpected genotype-phenotype association.

Our findings contribute to a growing awareness that protein polymorphism may often have phenotypic effects that are unrelated to the ‘canonical’ function of the protein in question. For example, genetic variation in enzymes of central metabolism can affect physiological phenotypes via mechanisms independent of pathway flux due to signalling functions of intermediary metabolites or nonenzymatic ‘moonlighting’ functions of the enzymes (Marden, 2013a, 2013b). The realization that there may be physiologically important auxiliary functions still waiting to be discovered in a protein as intensively studied as Hb highlights the importance of maintaining a wide field of vision when investigating causal connections between genotype and phenotype.

## METHODS

### Deer mouse populations and breeding designs

Wild adult deer mice were live-trapped at high altitude on the summit of Mount Evans (Clear Creek County, CO, USA; 4,350 m above sea level) (*P. m. rufinus*) and at low altitude on the Great Plains (Nine Mile Prairie, Lancaster County, NE, USA; 430 m above sea level) (*P. m. nebrascensis*). A highland male and lowland female were used to produce first-generation inter-population hybrids (F_1_), which were then raised to maturity and used for full-sibling matings to produce second-generation hybrid (F_2_) progeny. Each F_2_ hybrid was genotyped to determine the sequence of its α- and β-globin haplotypes, resulting in the 5 distinct combinations of highland and lowland haplotypes of α- and β-globin that were studied here: N=5 α^HH^β^HH^, N=5 α^HH^β^HL^, N=7 α^HH^β^LL^, N=4 α^LL^β^HH^ and N=5 α^LL^β^HL^. Other wild mice were bred in captivity to produce first-generation (G1) progeny within each population. Additional details on holding conditions, genotyping, and acclimations can be found in the Supplementary Methods.

### Physiological effects of haemoglobin genotype in inter-population hybrids

Acute hypoxia responses were measured during adulthood (1-1.5 years old) in unrestrained conditions in all F_2_ hybrids both before and after acclimation to hypobaric hypoxia (12 kPa O_2_ for 6 weeks), using barometric plethysmography, respirometry, and pulse oximetry techniques that we have described in previous studies (Ivy et al., 2020; Ivy and Scott, 2018, 2017). Individual mice were placed in a whole-body plethysmography chamber that was supplied with hyperoxic air (30 kPa O_2_, balance N_2_) and given 20-40 min to adjust. Mice were then maintained for 20 min at 30 kPa O_2_, after which they were exposed to 20 min stepwise reductions in PO_2_ of 21, 16, 12, 10, and 8 kPa. Breathing (total ventilation, breathing frequency, and tidal volume), rates of O_2_ consumption (V^˙^ O_2_), body temperature (T_b_), heart rate, and arterial O_2_ saturation were determined during the last 10 min at each PO_2_. Haematological measurements were also made before and after hypoxia acclimation, after at least 5 d recovery from measurements of acute hypoxia responses. Additional details of methods and calculations can be found in the Supplementary Methods.

### Physiological effects of manipulating Hb-O_2_ affinity with efaproxiral

Captive G_1_ populations of deer mice from high and low altitude, held in standard holding conditions in normoxia, were used to assess the acute effects of manipulating Hb-O_2_ binding.

Mice were placed in the plethysmography chamber and exposed to normoxic conditions (21 kPa O_2_) for 40 min to make baseline measurements. Mice were then removed and given an intraperitoneal injection of either saline or efaproxiral sodium at a volume of 20 ml per kg body mass (Fisher Scientific, Whitby, ON, Canada). Efaproxiral was prepared in sterile saline (0.9% NaCl solution) on the day of experiments and was administered at a dose of 200 mg per kg body mass. Mice were then returned to the chamber, and measurements were made for 50 min in normoxia and 20 min in hypoxia (12 kPa O_2_). Breathing, V^˙^ O_2_, T_b_, heart rate, and arterial O_2_ saturation were measured in the last 10 min of each exposure as described above. Every individual underwent both saline and efaproxiral injections, conducted in random order and separated by 1 week, and the efaproxiral dose used was determined in preliminary tests to have persistent effects on arterial O_2_ saturation for the duration of the experiment.

### Statistical analysis of physiological variables

Linear mixed-effects models were used in experiments with F_2_ hybrids to test for effects of α and β globin genotype, acclimation environment, and inspired PO_2_. They were also used in the efaproxiral experiments to test for effects of efaproxiral, mouse population, and inspired PO_2_. We initially tested for the random effects of sex and family, but they did not near statistical significance (P>0.10) and were therefore removed from the final models reported here. Holm-Šidák post-tests were used as appropriate. Statistical analysis was conducted using the lme4 package in R (v. 3.6.0) (Bates et al., 2015) with a significance level of P < 0.05. Values are reported as mean ± SEM.

### RNA-Seq library preparation and transcriptomic analysis

We used high-throughput sequencing (RNA-seq) (Wang et al., 2009) to test for effects of haemoglobin genotype on gene expression in the medulla of F_2_ hybrids sampled after hypoxia acclimation, using methods that have been previously described (Schweizer et al., 2019; Velotta et al., 2018) and are detailed in the Supplementary Methods. Raw sequences are deposited in the NCBI Short Read Archive (SRA accession PRJNA670858). Briefly, Illumina sequencing libraries were generated for 5 α^HH^β^HH^ mice and 5 α^LL^β^HL^ mice using TruSeq RNA Sample Preparation Kit v2 (Illumina, San Diego, CA, USA) and sequenced on an Illumina HiSeq2500 platform. After filtering, we obtained a total of 368.3 million reads with an average 36.8 million reads per individual (range = 33.3-44.5 million), and an average read length of approximately 120 bp. Reads were mapped to the *P. maniculatus bairdii* genome (Pman_1.0; GenBank accession: GCF_000500345.1), and transcript abundance was determined for a total of 16,082 genes that had at least 20 normalized reads per individual on average. We compared the level of transcript abundance between genotypes using whole-transcriptome differential expression analysis in *edgeR* (Robinson and Oshlack, 2010), along with weighted gene co-expression network analysis (WGCNA v. 1.41-1) (Langfelder and Horvath, 2008) to identify whether modules of co-expressed genes were differentially expressed between genotypes.

## ACKNOWLEDGEMENTS

This research was funded by a Natural Sciences and Engineering Research Council of Canada (NSERC) Discovery Grant to G.R.S (RGPIN-2018-05707). National Sciences Foundation (NSF) grants to Z.A.C. (IOS-1354934, IOS-1634219, IOS-1755411, and OIA-1736249) and J.F.S. (IOS-1354390 and OIA-1736249), and a National Institutes of Health (NIH) grant to J.F.S. (HL087216). Salary support was provided to C.M.I. by a NSERC Postgraduate Scholarship and an Ontario Graduate Scholarship, to O.H.W. by a NSERC Vanier Canada Graduate Scholarship, to R.M.S. by an NSF Postdoc Research Fellowship in Biology (1612859), to J.P.V. by a NIH National Heart, Lung and Blood Institute Research Service Award Fellowship (1F32HL136124-01), to S.C.C.-S. by a NSF (DBI) Postdoctoral Fellowship in Biology Award (#1612283), and to G.R.S. by the Canada Research Chairs Program.

## COMPETING INTERESTS

The authors declare no competing interests.

## DATA AVAILABILITY

Physiological data are deposited in Mendeley Data (DOI: 10.17632/mktd4vn3d7.1) and raw RNA-Seq sequences are deposited in the NCBI Short Read Archive (SRA accession PRJNA670858).

## AUTHOR CONTRIBUTIONS

G.R.S., J.F.S., and Z.A.C. designed the study. C.N., N.G.-P., J.P.V., and S.C.C.-S. carried out mouse breeding and genotyping. C.M.I. and O.H.W. ran and analyzed the *in vivo* experiments. C.M.I., C.N., R.M.S., J.P.V., E.E.P., and A.F. carried out tissue analyses and/or bioinformatics. C.M.I. and G.R.S. wrote the manuscript, and all authors edited the manuscript.

## SUPPLEMENTARY METHODS

### Deer mouse populations and breeding designs

Wild adult deer mice were live-trapped at high altitude on the summit of Mount Evans (Clear Creed County, CO, USA at 39°35’18”N, 105°38’38”W; 4,350 m above sea level) (*P. m. rufinus*) and at low altitude on the Great Plains (Nine Mile Prairie, Lancaster County, NE, USA at 40°52’12”N, 96°48’20.3”W; 430 m above sea level) (*P. m. nebrascensis*) and were transported to the University of Montana (elevation 978 m) or to McMaster University (elevation 50 m). The wild mice transported to Montana were used to produce one family of first-generation inter-population hybrids (F_1_), created by crossing a highland male and a lowland female. These F_1_ hybrids were raised to maturity and were used for full-sibling matings to produce 4 families of second-generation hybrid progeny (F_2_). These F_2_ hybrids (N=26) were raised to adulthood, a small volume of blood was obtained for genotyping (sampled from the facial vein and then stored at −80°C), and mice were then transported to McMaster for subsequent experiments (see below). These F_2_ hybrid mice were also used for a distinct study on aerobic capacity (Wearing et al., 2020). The wild mice transported to McMaster were bred in captivity to produce first-generation (G_1_) progeny within each population. All mice were held in a standard holding environment (∼23°C, 12 h:12 h light:dark photoperiod) under normal atmospheric conditions before experiments, and were provided with unlimited access to water and standard mouse chow. All animal protocols were approved by institutional animal research ethics boards.

Adult isoforms of tetrameric haemoglobin from *P. maniculatus* incorporate α-chain subunits that are encoded by two tandem gene duplicates, *HBA-T1* and *HBA-T2* (separated by 5.0 kb on Chromosome 8), and β-chain subunits that are encoded by two other tandem duplicates, *HBB-T1* and *HBB-T2* (separated by 16.2 kb on Chromosome 1) (Hoffmann et al., 2008; Natarajan et al., 2015a). We used a reverse-transcriptase PCR (RT-PCR) approach to obtain sequence data for all four of the adult-expressed α- and β-globin transcripts in the full panel of mice (Natarajan et al., 2015a; Storz et al., 2010). The RNeasy Plus Mini Kit (Qiagen, Valencia, CA, USA) was used to extract total RNA from red blood cells. We then amplified globin transcripts from 1 μg of extracted RNA using the One-Step RT-PCR system with Platinum *Taq* DNA polymerase High Fidelity (Invitrogen, Carlsbad, CA, USA). PCR cycling conditions were as follows: 1 cycle at 50 °C for 30 min, 1 cycle at 95°C for 15 min, 34 cycles at 94 °C for 30 s, 55 °C for 30 s, and 72 °C for 1 min, and then a final extension cycle at 72 °C for 3 min. For the α-globin transcripts, we used the same primer pair for *HBA-T1* and *HBA-T2* (forward: CTGATTCTCACAGACTCAGGAAG, reverse: CCAAGAGGTACAGGTGCGAG). For the β-globin transcripts, we used the same RT-PCR primer pair for *HBB-T1* and *HBB-T2* (forward: GACTTGCAACCTCAGAAACAGAC, reverse: GACCAAAGGCCTTCATCATTT). We cloned and sequenced the RT-PCR products using TOPO^®^ TA Cloning Kit (Life Technologies, Carlsbad, CA, USA), and we sequenced at least six clones per gene in order to recover both alleles from the paralogs. Full-length cDNAs of all expressed *HBA* and *HBB* genes were thereby sequenced at 6-fold coverage, and the haplotype phase of all variable sites was determined experimentally. We thus identified mice with the following combinations of lowland (L) and highland (H) haplotypes of α-globin and β-globin for use in subsequent experiments: N=5 α^HH^β^HH^, N=5 α^HH^β^HL^, N=7 α^HH^β^LL^, N=4 α^LL^β^HH^ and N=5 α^LL^β^HL^.

### Physiological measurements on F_2_ interpopulation hybrids

F_2_ hybrids were subjected to a series of measurements during adulthood (1-1.5 years old), both before and after exposure to chronic hypoxia. Acute hypoxia responses and haematology were first measured in mice held in normoxia. Four days later, the mice were moved into specifically designed hypobaric chambers that have been previously described (Ivy and Scott, 2017; Lui et al., 2015; McClelland et al., 1998) and were thus acclimated to hypoxia for 8 weeks (barometric pressure of 60 kPa, simulating the pressure at an elevation of 4,300 m; O_2_ pressure ∼12.5 kPa). During this time, mice were temporarily returned to normobaric conditions twice per week for <20 min for cage cleaning. Acute hypoxia responses were then measured again after 6-8 weeks in chronic hypoxia. Mice were finally euthanized after a full 8 weeks of hypoxia acclimation with an overdose of isoflurane followed by cervical dislocation, blood was collected for the second set of haematology measurements, and the medulla was sampled and stored at − 80°C for subsequent transcriptomic analyses (see below).

Acute hypoxia responses were measured in unrestrained mice using barometric plethysmography, respirometry, and pulse oximetry techniques that we have used in previous studies (Ivy et al., 2020; Ivy and Scott, 2018, 2017). Mice were placed in a whole-body plethysmography chamber (530 ml) that was supplied with hyperoxic air (30 kPa O_2_, balance N_2_) at 600 ml min^-1^. Mice were given 20-40 min to adjust to the chamber until relaxed and stable breathing and metabolism were observed. Mice were then maintained for an additional 20 min at 30 kPa O_2_, after which they were exposed to 20-min stepwise reductions in inspired O_2_ pressure (PO_2_) of 21, 16, 12, 10, and 8 kPa. Dry incurrent gases were mixed using precision flow meters (Sierra Instruments, Monterey, CA, USA) and a mass flow controller (MFC-4, Sable Systems, Las Vegas, NV, USA), such that the desired PO_2_ was delivered to the chamber at a constant flow rate of 600 ml min^-1^. At the end of this protocol, mice were removed from the chamber and returned to their home cage in the appropriate acclimation condition.

Breathing (total ventilation, breathing frequency, and tidal volume), rates of O_2_ consumption (V^˙^ O_2_), body temperature (T_b_), heart rate, and arterial O_2_ saturation were determined during the last 10 min at each PO_2_ as follows. Incurrent and excurrent air flows were subsampled at 200 ml min^-1^; incurrent air was continuously measured for O_2_ fraction (FC-10, Sable Systems), and excurrent air was analyzed for water vapour (RH-300, Sable Systems), dried with pre-baked drierite, and analyzed for O_2_ and CO_2_ fraction (FC-10 and CA-10, Sable Systems). These data were used to calculate V^˙^ O_2_, expressed in volumes at standard temperature and pressure (STP), using established equations (Lighton, 2008). Chamber temperature was continuously recorded with a thermocouple (TC-2000, Sable Systems). Breathing frequency and tidal volume were measured using the barometric method of whole-body plethysmography with established equations (Drorbaugh and Fenn, 1955; Jacky, 1980). Total ventilation was calculated as the product of breathing frequency and tidal volume. Total ventilation and tidal volume data are reported in volumes expressed at body temperature and pressure saturated (BTPS). Air convection requirement (ACR) is the quotient of total ventilation and V^˙^ O_2_. All of the above data was acquired using PowerLab 16/32 and Labchart 8 Pro software (ADInstruments, Colorado Springs, CO, USA). T_b_ was measured using thermosensitive passive transponders (micro LifeChips with Bio-therm technology; Destron Fearing, Dallas, TX, USA), which were implanted subdermally on the left side of the abdomen close to the leg ∼2 weeks before normoxic measurements were conducted, along with a hand-held scanner from the same manufacturer. Arterial O_2_ saturation and heart rate were measured using MouseOx Plus pulse oximeter collar sensors and data acquisition system (Starr Life Sciences, Oakmont, PA, USA). This was enabled by removing fur around the neck ∼2 days before experiments.

Blood was collected into heparinized capillary tubes for haematology, sampled from the facial vein under light anaesthesia (∼130 µl) for mice acclimated to normoxia, or by severing the jugular vein for mice that were euthanized and sampled after hypoxia acclimation. Blood Hb content was measured using Drabkin’s reagent according to the manufacturer’s instructions (Sigma-Aldrich, Oakville, ON, Canada). Haematocrit was measured by spinning the blood in the capillary tubes at 12,700 *g* for 5 min. Oxygen dissociation curves were generated at 37 °C for all mice using a Hemox Analyzer (TCS Scientific, New Hope, PA, USA) using 10 µl of whole blood in 5 ml of buffer (100 mmol l^-1^ HEPES, 50 mmol l^-1^ EDTA, 100 mmol l^-1^ KCl, 0.1% bovine serum albumin, and 0.2% antifoaming agent; TCS Scientific). Red blood cell O_2_ affinity (P_50_, the PO_2_ at which haemoglobin is 50% saturation with O_2_) was calculated using Hemox Analytic Software (TCS Scientific). These measurements of blood Hb content, haematocrit, and P_50_ have been previously published (Wearing et al., 2020) but are reported again here to provide insight into the measurements of acute hypoxia responses. In addition, the NO metabolites nitrite, S-nitrosothiols (SNO), and iron-nitrosyl and N-nitrosamine derivatives (FeNO + NNO) were measured by reductive chemiluminescence in plasma and red blood cells for a subset of hypoxia-acclimated mice following euthanasia, using a Sievers Nitric Oxide Analyzer (NOA model 280i, Boulder, CO, USA) and previously described protocols (Hansen and Jensen, 2010; Yang et al., 2003). Blood was sampled in dim light conditions, spun at 16,000 *g* for 2 min to separate plasma from red blood cells, which were then quickly frozen in liquid N_2_ and stored at − 80°C. Frozen samples (100 µl) were thawed and immediately incubated at room temperature for 5-10 min in the dark with a SNO-stabilizing solution (900 µl) described elsewhere (Yang et al., 2003) and then centrifuged (10,000 g for 2 min). The supernatant of each sample was injected into the NOA purge vessel in serial aliquots as is (300 µl, peak A) and after 2 min incubation with sulfanilamide (270 µl, peak B) and sulfanilamide and HgCl_2_ (270 µl, peak C) to obtain values for nitrite, SNO, and FeNO + NNO from the three peaks, as previously described (Yang et al., 2003).

### Physiology effects of inhibition of nitric oxide synthase

Based on the observed effects of α-globin genotype on both breathing pattern and blood NO metabolites, we used a pharmacological approach using CD1 lab-strain mice to investigate whether manipulating NO production affects breathing in normoxia acclimated mice. Ten mice (5 male, 5 female) were purchased from Charles River Laboratories and housed at McMaster University. Mice were held in normoxia in the same conditions described above for deer mice. To examine the effects of manipulating NO production *via* NO synthase (NOS), mice were first placed in the same plethysmograph as described above, and were exposed to normoxic conditions (21 kPa O_2_) for 30 min in order to make baseline measurements. Mice were then removed and given an intraperitoneal injection of either saline, N^ω^-nitro-L-arginine methyl ester hydrochloride (L-NAME, a universal NOS inhibitor; N5751, Sigma-Aldrich, Mississauga, ON, Canada), or S-methyl-L-thiocitrulline acetate salt (SMTC, a neuronal NOS inhibitor; M5171, Sigma-Aldrich) at a volume of 20 ml per kg body mass. L-NAME and SMTC were prepared in sterile saline (0.9% NaCl solution) on the day of experiments and were administered at a dose of 5 and 10 mg per kg body mass, respectively, based on previous experiments conducted in our lab and the literature (El Hasnaoui-Saadani et al., 2007; Pamenter et al., 2015; Pichon et al., 2009). Mice were returned to the chamber and measurements were made for 30 min in normoxia, followed by 30 min of hypoxia (12 kPa O_2_). Breathing, V^˙^ O_2_, and T_b_ were measured in the last 10 min of each exposure as described above. Every individual underwent all three injection treatments (saline, L-NAME, and SMTC), conducted in random order and separated by 2 days.

### Exome sequencing and linkage analysis

Based on the observed effects of α-globin on breathing pattern, we used an exome capture approach to determine genes that were linked to α-globin and might contribute to the differences in breathing frequency and tidal volume. Details on exome design, DNA capture, and high-throughput sequencing have been previously published (Schweizer et al., 2019). Briefly, we targeted the nuclear exome, 5000 randomly selected non-genic segments of 500 bp each, and promoter regions for a subset of genes. We extracted DNA from liver tissue, prepared genomic libraries using the NEBNext UltraII kit and with unique index sequences, following manufacturer’s protocols, then pooled libraries equimolarly. Using a Roche NimbleGen SeqCap EZ Library, we captured a total of 77,559,614 base pairs from 24 of 26 individuals. We performed target enrichment for the exome capture array and sequenced multiplexed samples within one lane of a HiSeq 4000 with 100 bp paired-end sequencing. To process sequence data and identify variants, we followed the recommendations of the Broad Institute GATK v3.7-0-gcfedb67 Best Practices pipeline (https://gatk.broadinstitute.org/).

After trimming sequence reads for a minimum base quality of 20 along with any remaining adapter sequence, we used *bwa mem* (Li and Durbin, 2010) to map them to the *Peromyscus maniculatus baiardii* genome (assembly Pman_1.0). We used a set of known variants (Schweizer et al., 2019) for GATK *Base Quality Score Recalibration*, then called novel variants using *HaplotypeCaller* and *GenotypeGVCFs*. We applied the following filters to the data: quality of depth <2.0, FS > 60, mapping quality < 40, mapping quality rank sum < −12.5, read position rank sum < −8.0, and individual filters of genotype quality > 20, minimum depth of coverage > 5, and called in at least 95% of individuals. Sequencing failed for one individual (only 0.58% of total reads for the lane, 1.35X mean depth over capture regions) and was excluded from further analysis. In the remaining 23 samples, overall sequencing quality was high, with a mean filtered depth of coverage over capture regions of 25.6X±5.3X, and a mean genotyping rate over 98% (Table S5). We thus detected and proceeded with a data set of 475,552 filtered bi-allelic SNPs (min. 95% call rate, Hardy-Weinberg Equilibrium filter <0.05, minor allele frequency <0.05, depth >5, genotype quality>20). We confirmed that the α-globin and β-globin genotypes in the exome data matched those obtained using the RT-PCR approach described above.

We next used the sequence data to determine and characterize the genes that were within the same linkage block as α-globin. We first determined the haplotype phase of all individuals for chromosome 8, the chromosome that contains α-globin. To ensure that we accurately characterized linkage along the chromosome, we re-processed our raw sequence reads, again following GATK Best Practices, but using the *P. maniculatus* version 2 reference sequence for chromosome 8 (rather than the v1 consisting of multiple contigs). We phased haplotypes using the software shapeit (O’Connell et al., 2014) with default parameters, except for the effective population size, which we obtained from demographic models for deer mice (Schweizer et al., 2019). We calculated linkage as the squared correlation coefficient r2 using the ‘--hap-r2’ option in vcftools. To effectively capture the linkage block, we modeled the decay of LD as a function of physical distance from the His50Pro mutation in α-globin. Following Storz et al. (Storz et al., 2012), we used nonlinear regression and a model of recombination-drift equilibrium with mutation (Hill and Weir, 1988), treating upstream and downstream of the His50Pro mutation separately. We subsequently characterized the region of chromosome 8 surrounding this mutation wherein the expected r2 value remained above 0.2 (Storz et al., 2012). The identified region extended ∼2.65 Mb upstream and ∼9 Mb downstream of the His50Pro mutation (Figure S6) and contained a total of 116 genes (Table S6), of which a subset of 43 genes (including α-globin genes) contained at least one missense SNP (Table S7). We then performed functional enrichment analysis to test for significantly enriched gene ontology (GO) terms among the linked genes. For each *P. maniculatus* gene, we identified the orthologous *Mus musculus* gene, then used gProfiler (Reimand et al., 2016, 2011, 2007) to analyze GO term enrichment. We used strong hierarchical filtering (returning only the best term per parent term) to identify enriched gene functional categories above a false discovery rate corrected significance of 0.05. GO enrichment analysis performed for all 116 genes within the identified region of linkage identified a number of significant categories, most notably terms related to gamma-aminobutyric acid (GABA) signaling (Table S8). However, when GO enrichment analysis was performed for only the 43 linked genes that contained a missense SNP, the only significant categories that remained were “hemoglobin complex” (GO:0005833; P_adj_ = 0.0126), “oxygen binding” (GO:0019825; P_adj_ = 0.0178), and 3 serine hydrolase activity categories (GO:0017171, 0008236, 0004252; P_adj_ = 0.0178 for all). Running the latter analysis without the 2 duplicate α-globin genes removed the “hemoglobin complex” and “oxygen binding” categories, such that only categories related to serine hydrolase activity were enriched among the linked genes with missense SNPs.

### RNA-Seq library preparation and transcriptomic analysis

We used high-throughput sequencing (RNA-seq) (Wang et al., 2009) to test for the effects of haemoglobin genotype on gene expression in the medulla of F_2_ hybrids, from 5 α^HH^β^HH^ mice and 5 α^LL^β^HL^ mice sampled after hypoxia acclimation. We powdered the medulla under liquid N_2_ and extracted total RNA from 15 mg of tissue using TRI Reagent (Sigma-Aldrich, St. Louis, MO, USA), and then assessed RNA quality using TapeStation (RIN > 7; Agilent Technologies, Santa Clara, CA, USA). We generated Illumina sequencing libraries using TruSeq RNA Sample Preparation Kit v2 (Illumina, San Diego, CA, USA) beginning with 1 µg of RNA. The libraries were sequenced as 100 nt single-end reads on an Illumina HiSeq2500 platform, with all 10 individuals in one lane using Illumina index adaptors. We performed a series of filtering steps to remove artifacts generated during sequencing. Reads with an average Phred quality score of less than 30 were removed from each library, after which low-quality bases were removed from the remaining sequences (Trimmomatic function for Illumina sequence data in Galaxy (Bolger et al., 2014)). Adaptor sequences were trimmed from sequence reads. These quality control steps yielded a total of 368.3 million reads with an average 36.8 million reads per individual (range = 33.3-44.5 million), and an average read length of approximately 120 bp. Raw sequences are deposited in the NCBI Short Read Archive (SRA accession PRJNA670858).

Reads for each individual were mapped to the *P. maniculatus bairdii* genome (Pman_1.0; GenBank accession: GCF_000500345.1) using bwa (Li and Durbin, 2010), and we then computed transcript abundance values in Galaxy. We used *featureCounts* (Liao et al., 2014) to generate a table of transcript abundances. Sequence reads mapped to a total of 28,722 *P. maniculatus* genes. Since genes with low read counts are subject to increased measurement error (Robinson and Smyth, 2007) we excluded those with less than an average of 20 normalized reads per individual, leaving a total of 16,082 genes.

We compared the level of transcript abundance between α^HH^β^HH^ and α^LL^β^HL^ genotypes using whole-transcriptome differential expression analysis in *edgeR* (Robinson and Oshlack, 2010). The function *calcNormFactors* was used to normalize read counts across libraries, after model dispersion was estimated for each transcript separately using the function *estimateDisp* (McCarthy et al., 2012). We tested for differences in transcript abundance by first fitting a quasi-likelihood negative binomial generalized linear model to raw count data (*glmQLFit* function), which included a single main effect of genotype. P-values were calculated using a quasi-likelihood F test using the *glmGLFTest* function. We controlled for multiple testing by enforcing a transcriptome-wide false discovery rate correction of 0.05 (Benjamini and Hochberg, 1995).

Weighted gene co-expression network analysis (WGCNA v. 1.41-1) (Langfelder and Horvath, 2008) was used to identify modules of co-expressed genes and to then examine whether module expression differed between genotypes. Raw read counts were normalized by total library size and log-transformed using the *edgeR* functions *calcNormFactors* and *cpm*, respectively (Robinson et al., 2010). Module detection was performed using the *blockwiseModules* function in WGCNA with default parameters (Langfelder and Horvath, 2008). This involved calculating Pearson correlations of transcript abundance between pairs of genes, with an adjacency matrix computed by raising the correlation matrix to a soft thresholding power of β = 7. Soft thresholding was performed to achieve an approximately scale-free topology, an approach that favours strong correlations over weak ones (Zhang and Horvath, 2005). A β of 7 was chosen because it represents the value where improvement of scale-free topology model fit begins to decrease with increasing threshold of power. A topological overlap measure was computed from the resulting adjacency matrix for each gene pair. Topologically based dissimilarity was then calculated and used as input for average linkage hierarchical clustering in the creation of cluster dendrograms for the medulla. Modules were identified as branches of the resulting cluster tree using the dynamic tree-cutting method (Langfelder and Horvath, 2008).

WGCNA identified clusters of genes with highly correlated expression profiles across all 10 transcriptomes that were considered to be modules of co-expressed genes. ANOVA on rank-transformed module eigengene values was then used to test for the effects of α-globin genotype across modules, with P-values for the pairwise comparisons within a module corrected for multiple testing using a false discovery rate of 0.05 (Plachetzki et al., 2014).

### Globin detection in the brainstem using Western Blots

A distinct group of mice from captive G_1_ populations from high- and low-altitude were used to measure globin proteins in the medulla and pons. Mice (n=4 from each population) were acclimated to hypobaric hypoxia as described above for 8 weeks. Mice were then euthanized with an overdose of isoflurane, the chest was quickly opened to reveal the heart, an incision was made on the right atrium, and mice were perfused via the left ventricle with phosphate-buffered saline (PBS; 137 mmol l^-1^ NaCl, 2.68 mmol l^-1^ KCl, 10.0 mmol l^-1^ Na_2_HPO_4_, 1.76 mmol l^-1^ KH_2_PO_4_) to flush blood from the circulation of the brainstem. The medulla and pons were then isolated, sampled, and stored at −80°C.

Western blot analysis was carried out according to (Towbin et al., 1979) to assess α and β globin protein expression in the medulla and pons. Medulla and pons tissues were homogenized in ice-cold radioimmunoprecipitation assay buffer (RIPA) containing 1 mM phenylmethylsulfonyl fluoride and 1X Protein Stabilizing Cocktail (ThermoFisher Scientific, Cat: 89806, Waltham, MA, USA). The homogenate was centrifuged at 13,000 rpm for 10 min at 4°C and the protein content of the supernatant was quantified using the Bradford assay (following instructions from the manufacturer, BioRad, Hercules, CA, USA). These protein isolates (∼25 µg) were separated using 15% SDS-PAGE, along with a positive control (∼25 µg protein isolated from deer mouse blood hemolysate) and a pre-stained protein marker (PageRuler^TM^ Prestained Protein Ladder 10 to 180 kDa, cat# 26616, ThermoFisher Scientific). The proteins were transferred to nitrocellulose membranes (0.2 µm pore size) at 4 °C overnight using a Mini Trans-Blot cell (Bio-Rad, Hercules, CA, USA) at 30 V. The membrane was blocked by incubating in TBST buffer (20 mM Tris, 150 mM NaCl, 0.1% Tween 20) containing 5% non-fat milk powder for 1 h at room temperature. The membrane was washed three times with TBST for 10 min each. The membrane was then incubated overnight at 4 °C in TBST containing 5% BSA, anti-GAPDH primary antibody as a loading control (1:10000 dilution, ab181603, Abcam, Cambridge, MA, USA), and one of the anti-globin primary antibodies: anti-haemoglobin subunit alpha (1:200, ab92492, Abcam) or anti-haemoglobin subunit beta (1:1000, ab214049, Abcam). The membrane was then washed three times with TBST for 10 min each, and incubated for 2 h at room temperature in TBST containing 5% BSA and secondary antibody (goat anti-rabbit IgG (HRP), 1:2000, ab205718, Abcam). The membrane was again washed three times with TBST for 10 min each, and then developed in 10 ml of 50 mM Tris (pH 7.6) containing 6 mg of 3,3’-diaminobenzidine (DAB) and 10 µl of hydrogen peroxide. Developing time ranged from 3 to 10 min, dependent on the band intensity of the house-keeping protein (GADPH) and the background of the membrane.

## SUPPLEMENTARY FIGURES AND TABLES

**Figure S1.**
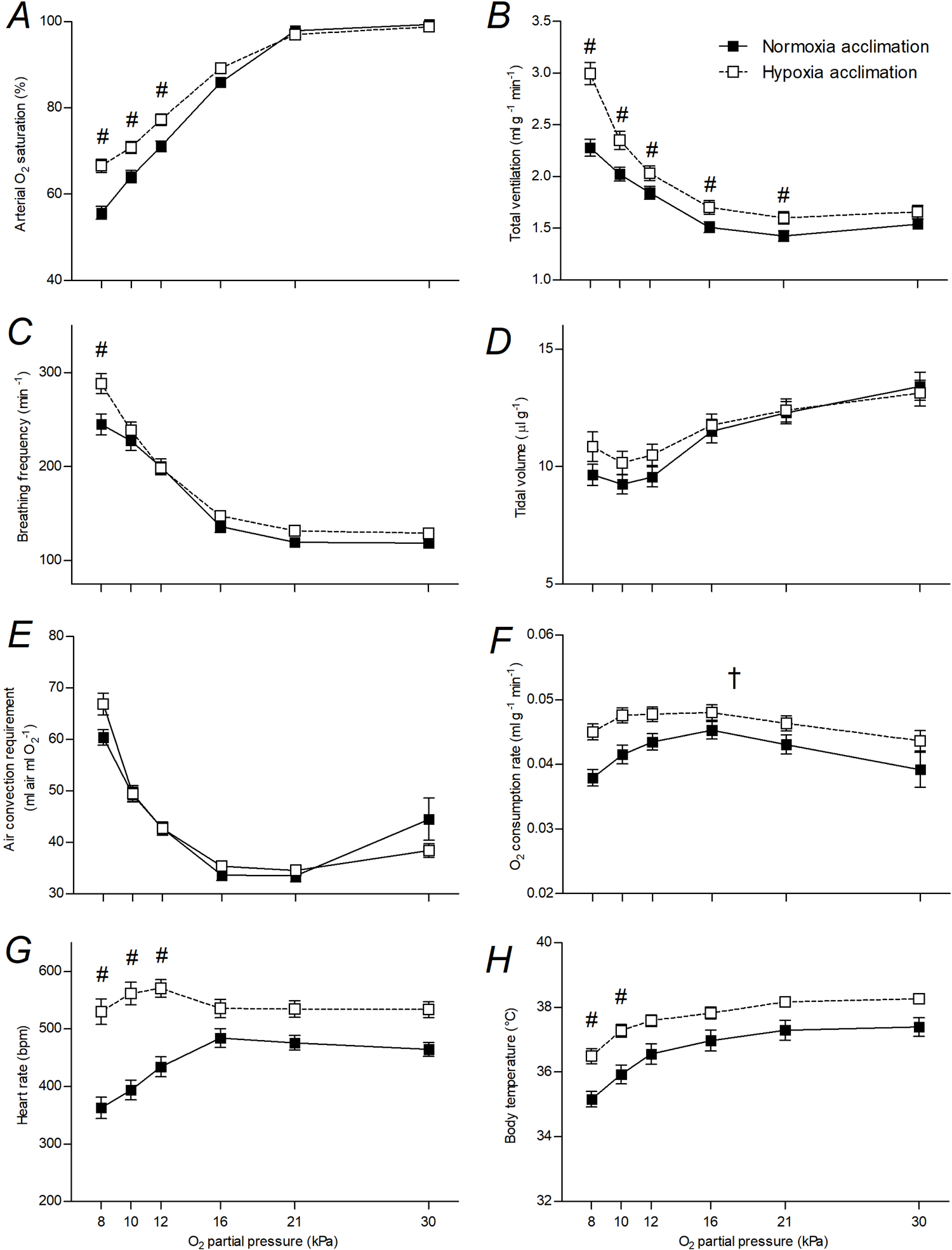
Chronic exposure to hypoxia affected the responses to acute hypoxia in F_2_ hybrid deer mice. Values are mean ± SEM. # represents a significant pairwise difference between acclimation groups within a PO_2_ using Holm-Šidák post-tests (N=26).

**Figure S2.**
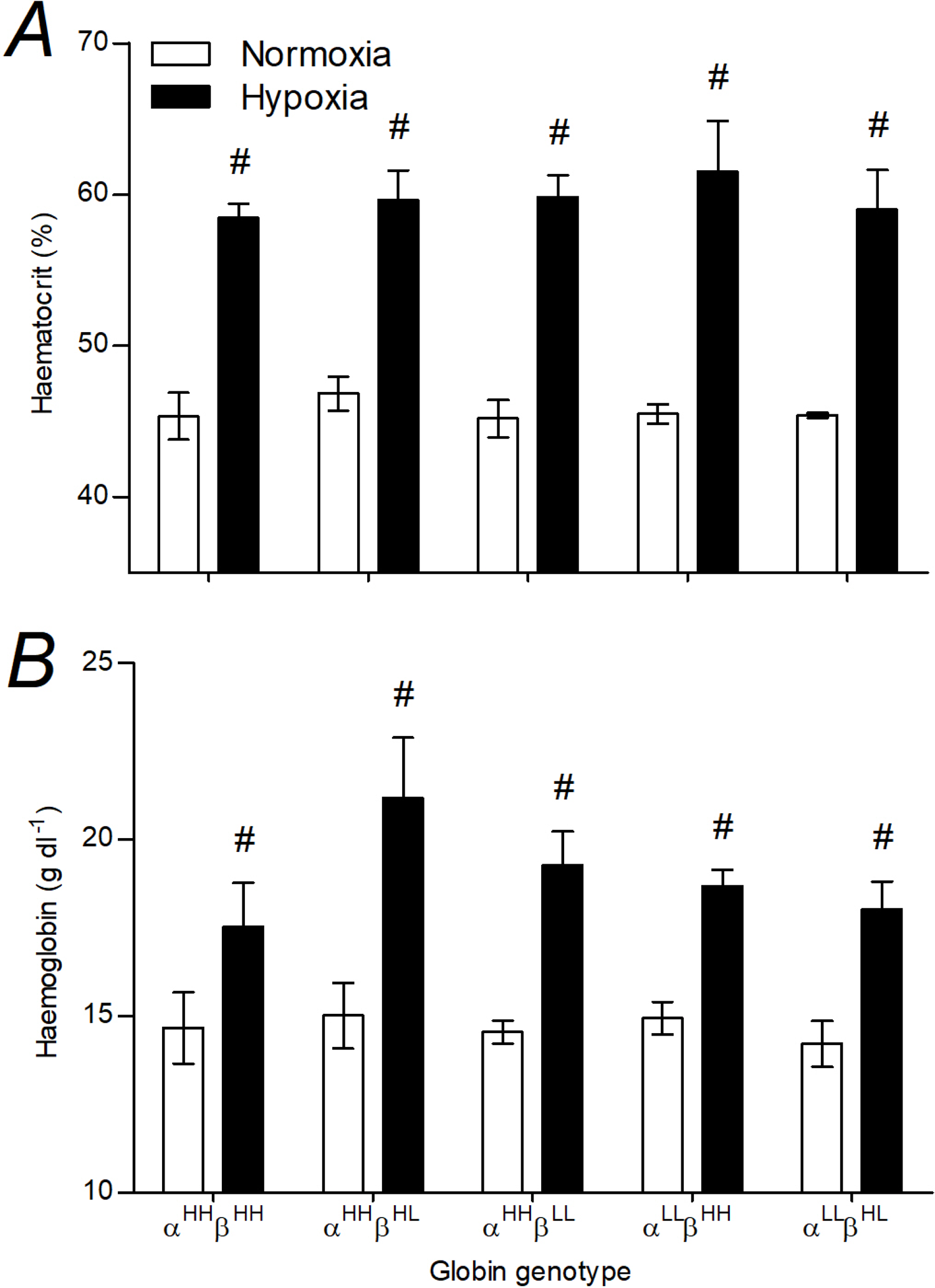
Haematocrit (A) and whole-blood haemoglobin content (B) were not influenced by α- or β-globin genotype in F_2_ hybrid deer mice, but increased after hypoxia acclimation. α^HH^β^HH^ represents mice that are homozygous for the highland α-globin and β-globin genotype (N=5), α^HH^β^HL^ represents mice that are homozygous for the highland α-globin genotype and heterozygous β-globin genotype (N=5), α^HH^β^LL^ represents mice that are homozygous for the highland α-globin genotype and homozygous lowland for the β-globin genotype (N=7), α^LL^β^HH^ represents mice that are homozygous for the lowland α-globin genotype and homozygous for the highland β-globin genotype (N=4), α^LL^β^HL^ represents mice that are homozygous for the lowland α-globin genotype and heterozygous β-globin genotype. Values are mean ± SEM. # denotes significant pairwise differences using Holm-Šidák post-tests between acclimation environments within a genotype.

**Figure S3.**
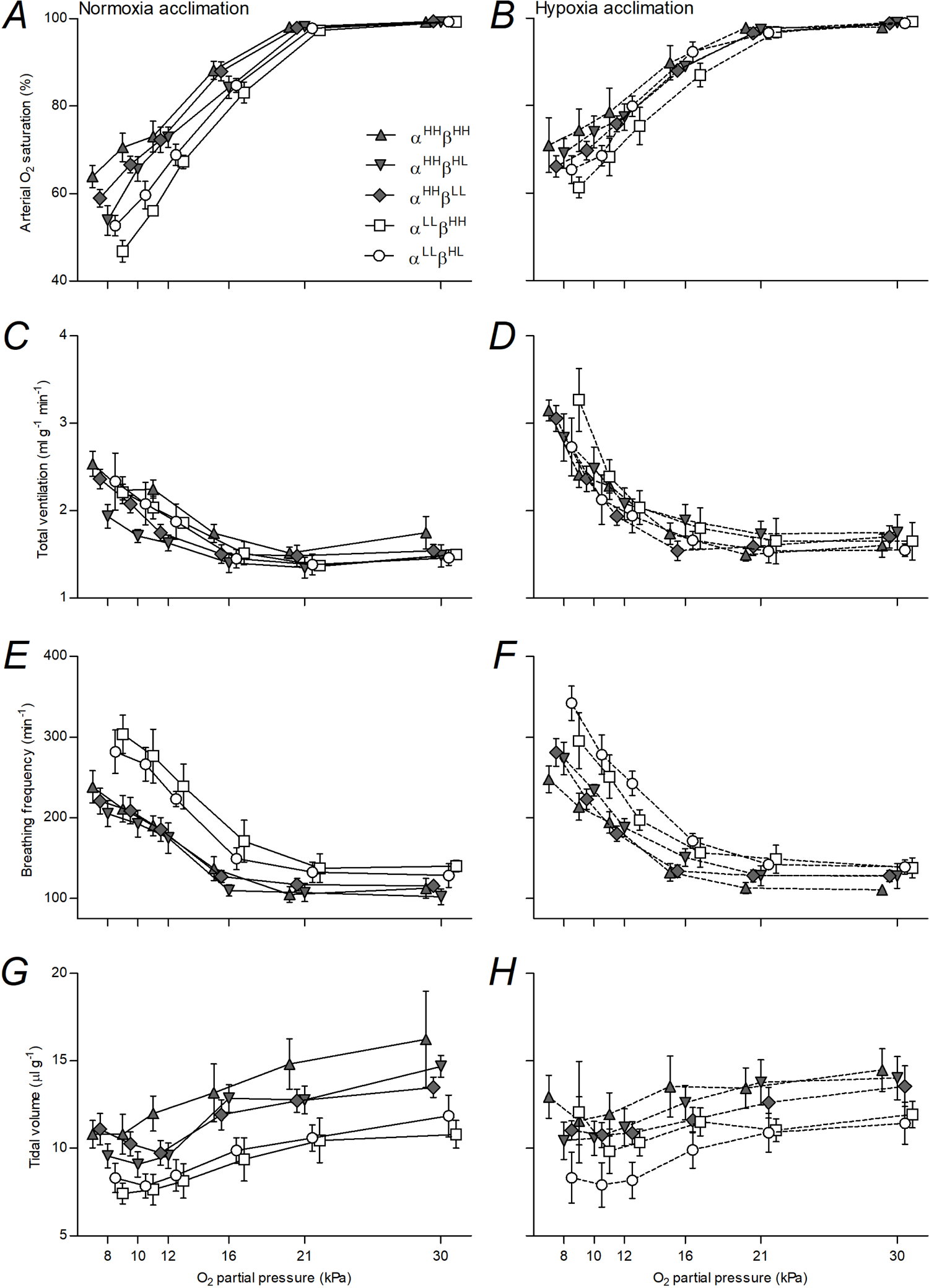
Arterial O_2_ saturation (A,B), total ventilation (C,D), breathing frequency (E,F), and tidal volume (G,H) responses of F_2_ hybrid mice before (A,C,E,G) and after (B,D,F,H) hypoxia acclimation in F_2_ hybrid mice with different α- and β-globin haplotypes. Values are mean ± SEM (genotypes and N as in Figure S2), symbols at each O_2_ partial pressure are offset for clarity.

**Figure S4.**
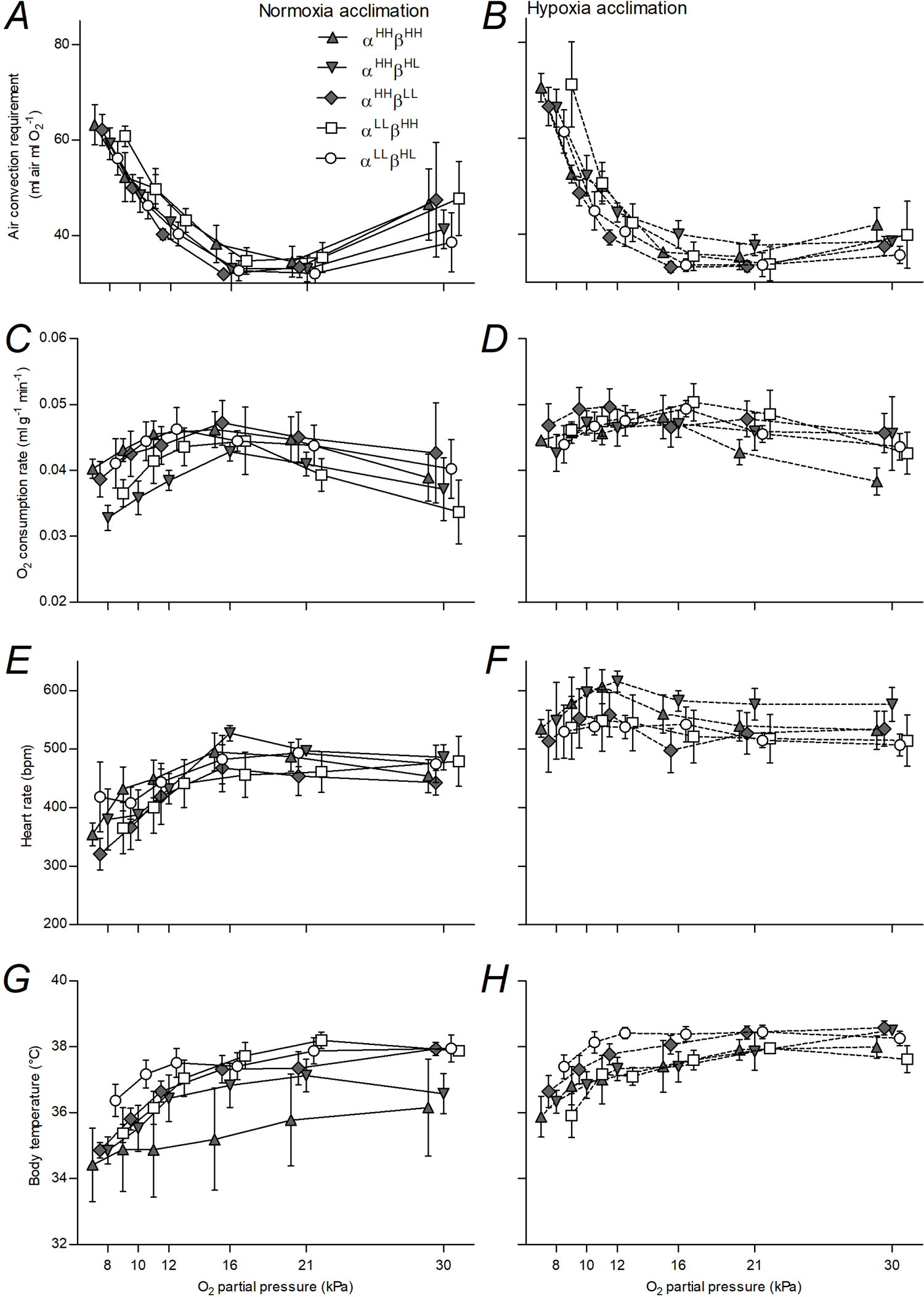
Air convection requirement (A,B), O_2_ consumption rate (C,D), heart rate (E,F), and body temperature (G,H) responses of F_2_ hybrid mice before (A,C,E,G) and after (B,D,F,H) hypoxia acclimation in F_2_ hybrid mice with different α- and β-globin haplotypes. Values are mean ± SEM (genotypes and N as in Figure S2), symbols at each O_2_ partial pressure are offset for clarity.

**Figure S5.**
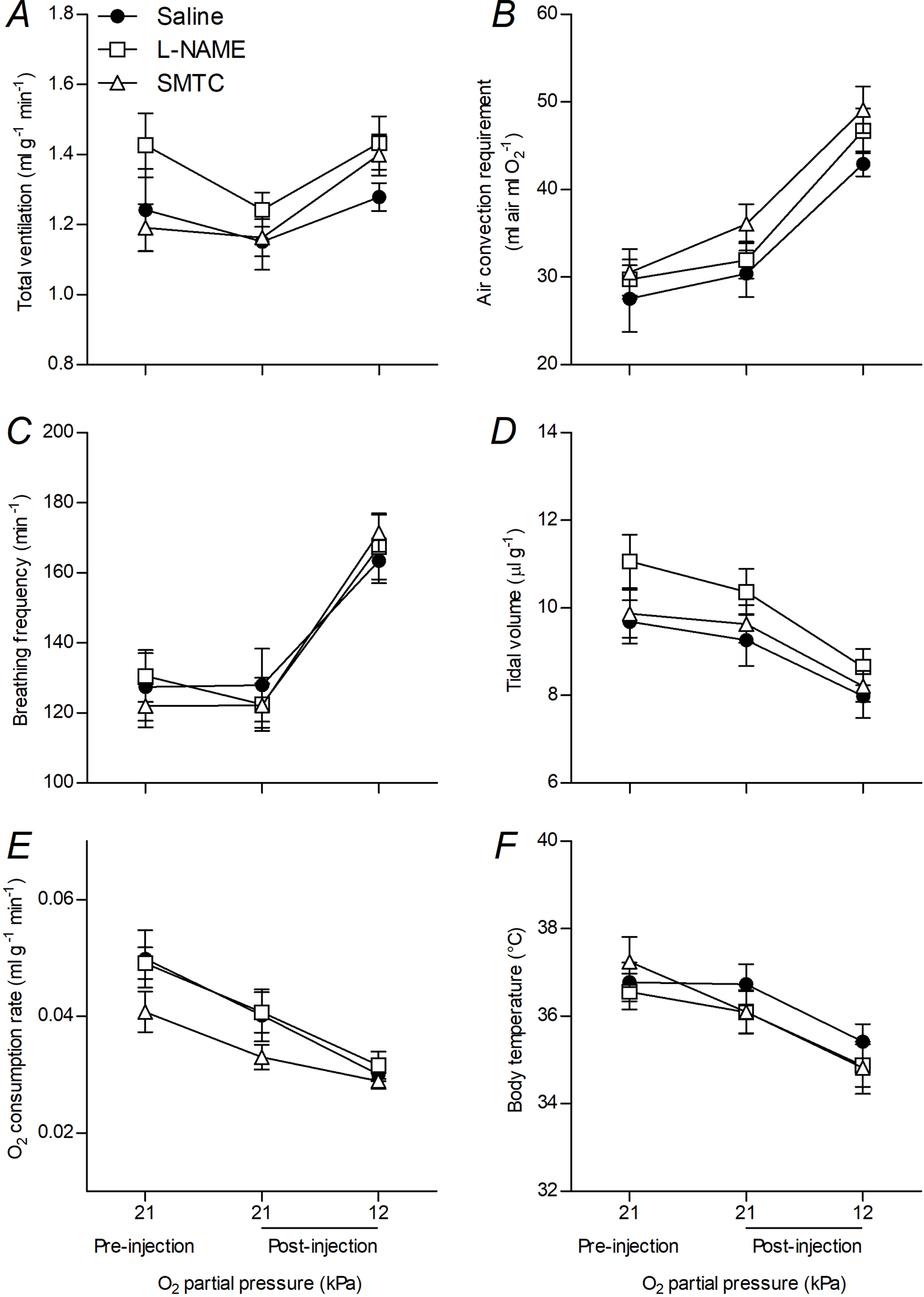
Nitric oxide synthase inhibition with L-NAME or SMTC did not influence total ventilation (A), air convection requirement (B), breathing frequency (C), tidal volume (D), O_2_ consumption rate (E), or body temperature (F) in lab-strain CD1 mice in normoxia or after acute exposure to hypoxia (12 kPa O_2_). Values are mean ± SEM (N=10 for each treatment).

**Figure S6.**
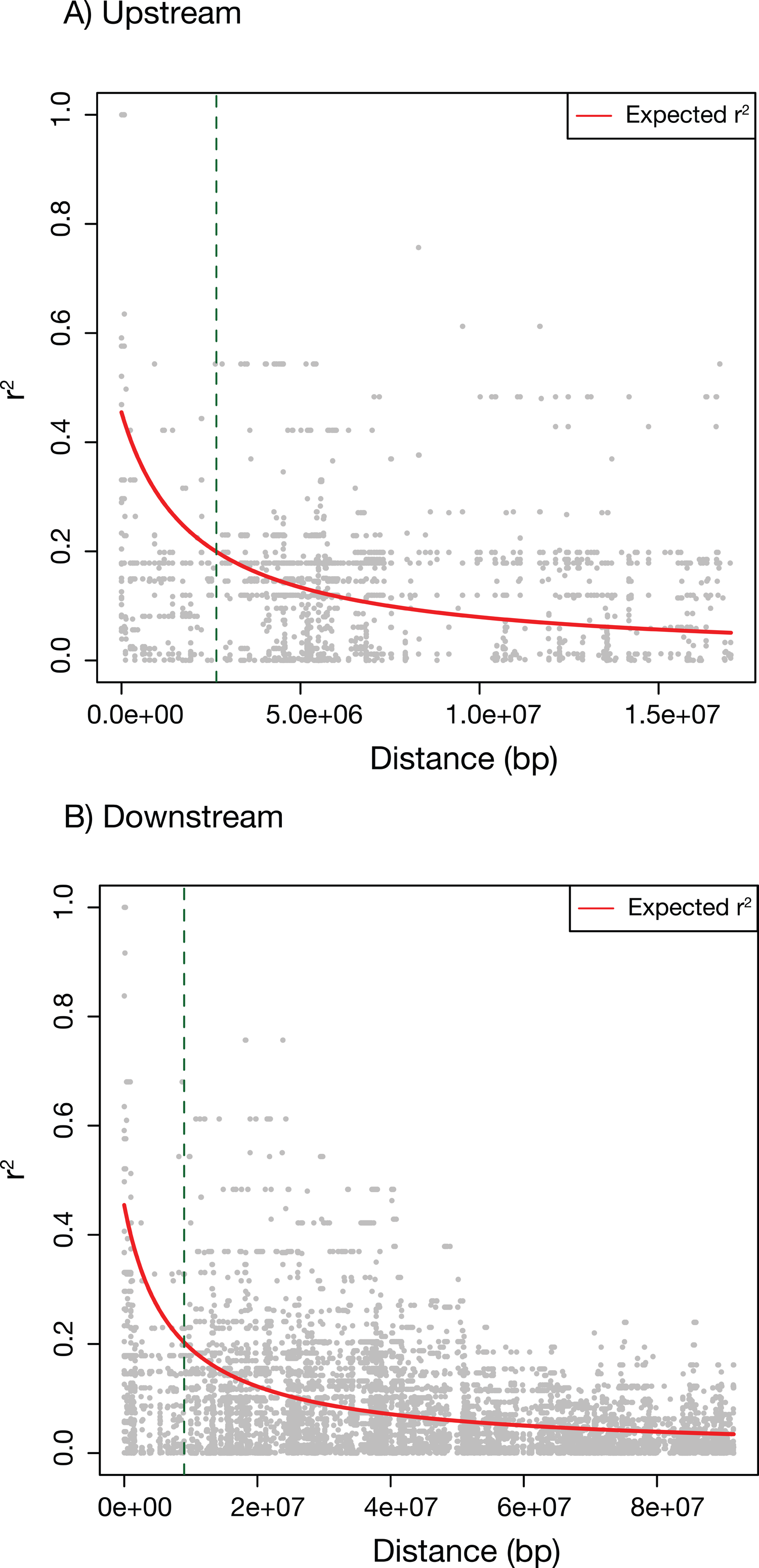
Rate of LD decay (A) upstream and (B) downstream of the His50Pro mutation in α-globin. In each plot, gray symbols indicate the r^2^ value for individual SNPs, while the red line shows the fitted nonlinear regression of r^2^ against physical distance using the Hill and Weir (1988) mutation-recombination-drift model. Vertical green dashed lines are the physical position where the fitted line falls below a r^2^ of 0.2.

**Figure S7.**
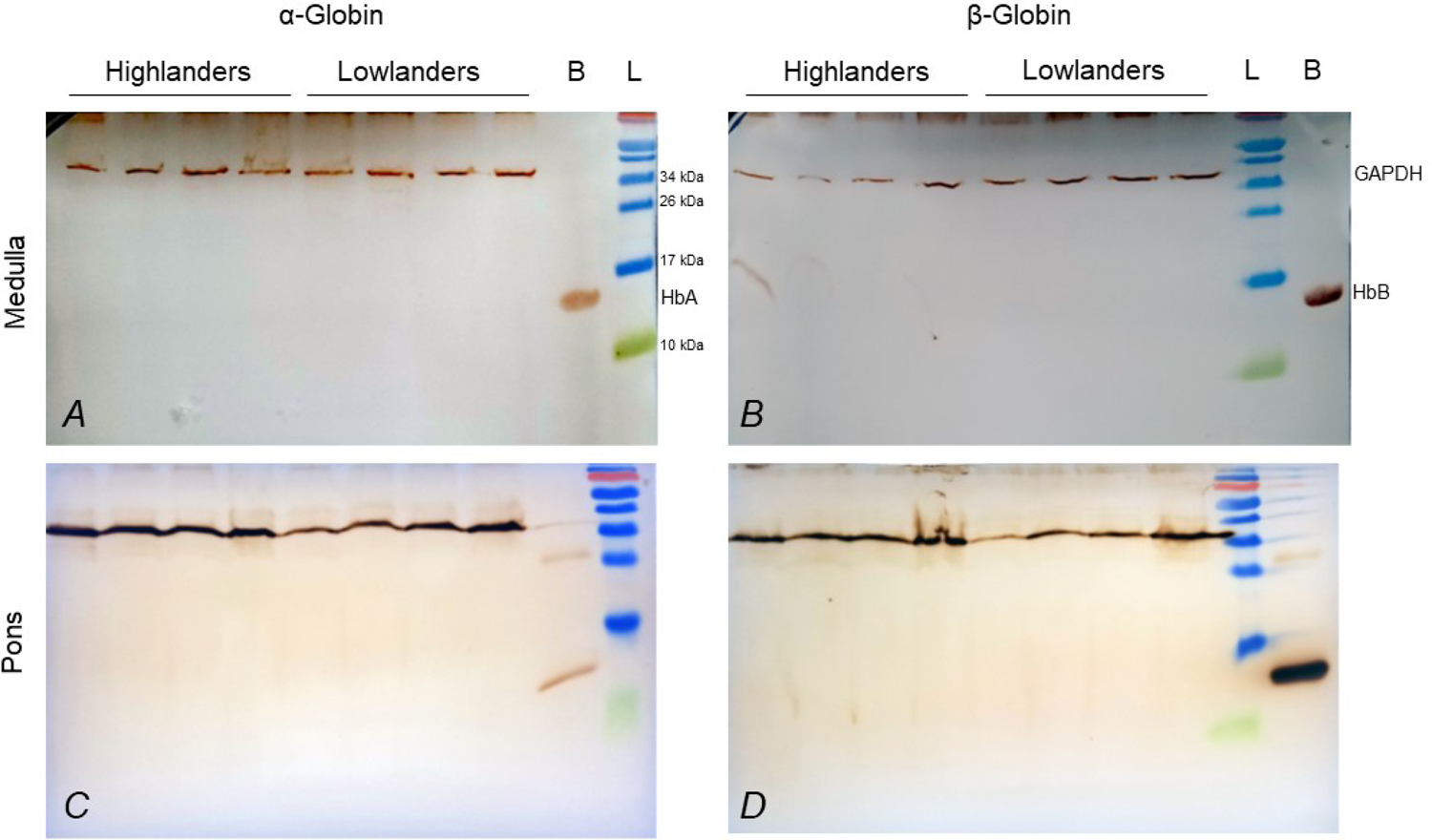
Western blots for α-globin (A,C) or β-globin (B,D) and for glyceraldehyde 3-phosphate dehydrogenase (GADPH; loading control) in protein isolates from medulla (A,B) and pons (C,D) tissues of high- and low-altitude populations of deer mice. B, deer mice blood hemolysate (positive control); L, protein ladder; HbA, α-globin; HbB, β-globin.

**Table S1.** Results of linear mixed-effects models of physiological responses to acute stepwise hypoxia in F_2_ hybrid deer mice

| | | $\alpha$ | $\beta$ | Env | PiO<sub>2</sub> | α*Env | β*Env | Env*PO<sub>2</sub> | α*PO<sub>2</sub> | β*PO<sub>2</sub> |
| --- | --- | --- | --- | --- | --- | --- | --- | --- | --- | --- |
| Arterial O<sub>2</sub> saturation | F | 6.9202 | 0.3438 | 15.406 | 579.58 | 0.6002 | 0.5098 | 4.6531 | n.s. | n.s. |
|  | P | **0.0115** | 0.7109 | **0.0003** | **<0.0001** | 0.5531 | 0.4789 | **0.0004** | n.s. | n.s. |
| Total ventilation | F | 0.2883 | 1.2737 | 12.333 | 215.46 | 0.1507 | 0.2229 | 13.5741 | n.s. | 2.4262 |
|  | P | 0.5938 | 0.2893 | **0.0009** | **<0.0001** | 0.6997 | 0.8011 | **<0.0001** | n.s. | **0.0091** |
| Breathing frequency | F | 20.612 | 0.1032 | 4.0450 | 346.65 | 0.5260 | 1.6550 | 5.1967 | 6.2340 | n.s. |
|  | P | **<0.0001** | 0.9021 | **0.0501** | **<0.0001** | 0.4721 | 0.2028 | **0.0001** | **<0.0001** | n.s. |
| Tidal volume | F | 15.581 | 1.3451 | 0.8357 | 61.040 | 0.1373 | 0.1466 | n.s. | n.s. | n.s. |
|  | P | **0.0003** | 0.2703 | 0.3653 | **<0.0001** | 0.7128 | 0.8640 | n.s. | n.s. | n.s. |
| Air convection requirement | F | 2.0682 | 47.003 | 0.1411 | 118.69 | 0.1417 | 0.3870 | 3.9349 | n.s. | n.s. |
|  | P | 0.1570 | 0.1087 | 0.7088 | **<0.0001** | 0.7084 | 0.6814 | **0.0019** | n.s. | n.s. |
| O<sub>2</sub> consumption rate | F | 0.5474 | 0.8862 | 7.4953 | 10.342 | 0.0001 | 0.0364 | n.s. | n.s. | n.s. |
|  | P | 0.4630 | 0.4190 | **0.0087** | **<0.0001** | 0.9991 | 0.9643 | n.s. | n.s. | n.s. |
| Heart rate | F | 1.0164 | 1.5240 | 33.048 | 10.583 | 0.6605 | 0.0057 | 13.524 | n.s. | n.s. |
|  | P | 0.3185 | 0.2284 | **<0.0001** | **<0.0001** | 0.4208 | 0.9943 | **<0.0001** | n.s. | n.s. |
| Body temperature | F | 7.8256 | 3.5823 | 13.469 | 76.840 | 2.1004 | 0.1326 | 1.9356 | n.s. | n.s. |
|  | P | **0.0074** | **0.0365** | **0.0006** | **<0.0001** | 0.1543 | 0.8762 | 0.0890 | n.s. | n.s. |

Env, acclimation environment, PiO<sub>2</sub>, O<sub>2</sub> partial pressure of inspired air during acute stepwise hypoxia; n.s. not significant

**Table S2.**
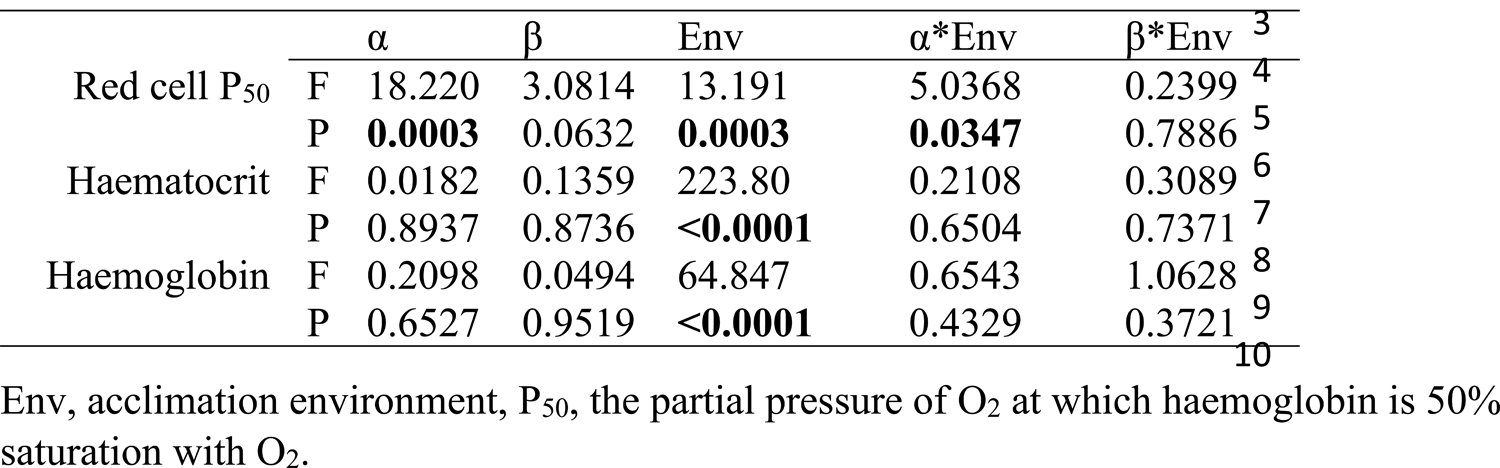
Results of linear mixed-effects models of blood responses to chronic hypoxia in F_2_ hybrid deer mice Env, acclimation environment, P_50_, the partial pressure of O_2_ at which haemoglobin is 50% saturation with O_2_.

**Table S3.** Nitric oxide (NO) metabolites measured in plasma and red blood cells after hypoxia acclimation were altered by α-globin genotype in F_2_ hybrid deer mice.

| | $\alpha^{LL}$ | $\alpha^{HH}$ | P |
| --- | --- | --- | --- |
| Plasma |  |  |  |
| Nitrite | 1.289 ± 0.255 (3) | 0.742 ± 0.070* (6) | **0.027** |
| SNO | 0.307 ± 0.045 (3) | 0.401 ± 0.061 (6) | 0.467 |
| Red blood cells |  |  |  |
| Nitrite | 0.270 ± 0.151 (5) | 0.864 ± 0.231 (8) | 0.089 |
| SNO | 0.484 ± 0.154 (4) | 0.485 ± 0.040 (8) | 0.994 |
| FeNO+NNO | 0.401 ± 0.186 (5) | 0.157 ± 0.054 (8) | 0.152 |

Values are mean ± SEM (n) in units $\mu\text{mol l}^{-1}$ ; SNO, S-nitroso compounds (SNO); FeNO+NNO, iron-nitrosyl and N-nitroso compounds. Asterisks indicate a significant difference between genotypes (t-test, $P < 0.05$ ).

**Table S4.** Results of linear mixed-effects models of the effects of nitric oxide inhibition and inspired PO2 in CD1 mice

| Variable |  | Treatment | PO<sub>2</sub> | Treatment*PO<sub>2</sub> | Random effect of sex |
| --- | --- | --- | --- | --- | --- |
| Total ventilation | F | 1.705 | 6.461 | 1.143 | - |
|  | P | 0.201 | **0.003** | 0.346 | n.s. |
| Air convection | F | 2.190 | 79.58 | 0.701 | - |
| requirement | P | 0.132 | **<0.0001** | 0.595 | **0.009** |
| Breathing frequency | F | 0.083 | 44.86 | 0.522 | - |
|  | P | 0.921 | **<0.0001** | 0.720 | n.s. |
| Tidal volume | F | 1.173 | 36.95 | 1.262 | - |
|  | P | 0.325 | **<0.0001** | 0.296 | n.s. |
| Oxygen consumption | F | 3.938 | 29.80 | 0.692 | - |
|  | P | **0.032** | **<0.0001** | 0.601 | **0.001** |
| Body temperature | F | 0.307 | 30.99 | 1.179 | - |
|  | P | 0.738 | **<0.001** | 0.330 | n.s. |

PO<sub>2</sub>, partial pressure of O<sub>2</sub>; n.s., not significant

Table S5 Summary of sequencing results for 24 individuals captured and sequenced with custom exome. Note that individual HL_F2_01_4r had poor sequencing coverage and was excluded from further analyses.

Table S6. Genes located within the ∼11.65 Mb linked region containing α-globin.

Table S7. Missense SNPs located within the ∼11.65 Mb linked region containing α-globin.

Table S8. GO enrichment results for the 116 genes located within the ∼11.65 Mb linked region containing α-globin.

**Table S9.** Results of linear mixed-effects models of the effects of efaproxiral and inspired PO_2_ in highland and lowland populations of deer mice

|  |  | Pop | Treat | PO<sub>2</sub> | Pop*PO<sub>2</sub> | Treat*PO<sub>2</sub> | Pop*Treat | Pop*Treat*PO<sub>2</sub> |
| --- | --- | --- | --- | --- | --- | --- | --- | --- |
| Arterial O<sub>2</sub> saturation | F | 4.1686 | 39.146 | 569.12 | 4.3395 | 11.410 | 10.715 | 3.2306 |
|  | P | 0.0684 | **<0.0001** | **<0.0001** | **0.0183** | **<0.0001** | **0.0019** | **0.0479** |
| Breathing frequency | F | 17.898 | 1.1134 | 39.657 | n.s | n.s | n.s | n.s |
|  | P | **0.0017** | 0.2958 | **<0.0001** | n.s | n.s | n.s | n.s |
| Tidal volume | F | 0.0460 | 0.5733 | 6.9964 | n.s | n.s | 7.8920 | n.s |
|  | P | 0.8344 | 0.4521 | **0.0019** | n.s | n.s | **0.0068** | n.s |
| Total ventilation | F | 5.6065 | 0.0175 | 11.034 | n.s | n.s | 7.8799 | n.s |
|  | P | **0.0394** | 0.8953 | **<0.0001** | n.s | n.s | **0.0069** | n.s |
| Air convection | F | 1.0930 | 0.6065 | 15.154 | n.s | n.s | n.s | n.s |
| requirement | P | 0.3204 | 0.4393 | **<0.0001** | n.s | n.s | n.s | n.s |
| O<sub>2</sub> consumption rate | F | 26.982 | 0.288 | 4.3785 | n.s | n.s | 7.8473 | n.s |
|  | P | **0.0004** | 0.5931 | **0.0171** | n.s | n.s | **0.0070** | n.s |
| Heart rate | F | 2.5305 | 2.5382 | 9.5632 | n.s | 5.7649 | 6.7673 | n.s |
|  | P | 0.1427 | 0.1170 | **0.0003** | n.s | **0.0054** | **0.0119** | n.s |
| Body temperature | F | 0.0024 | 1.8942 | 5.6830 | n.s | n.s | 12.483 | n.s |
|  | P | 0.9619 | 0.1742 | **0.0057** | n.s | n.s | **0.0008** | n.s |

Pop, population; Treat, treatment (saline or efaproxiral); PO<sub>2</sub>, partial pressure of O<sub>2</sub>; n.s., not significant

**Table S10.**
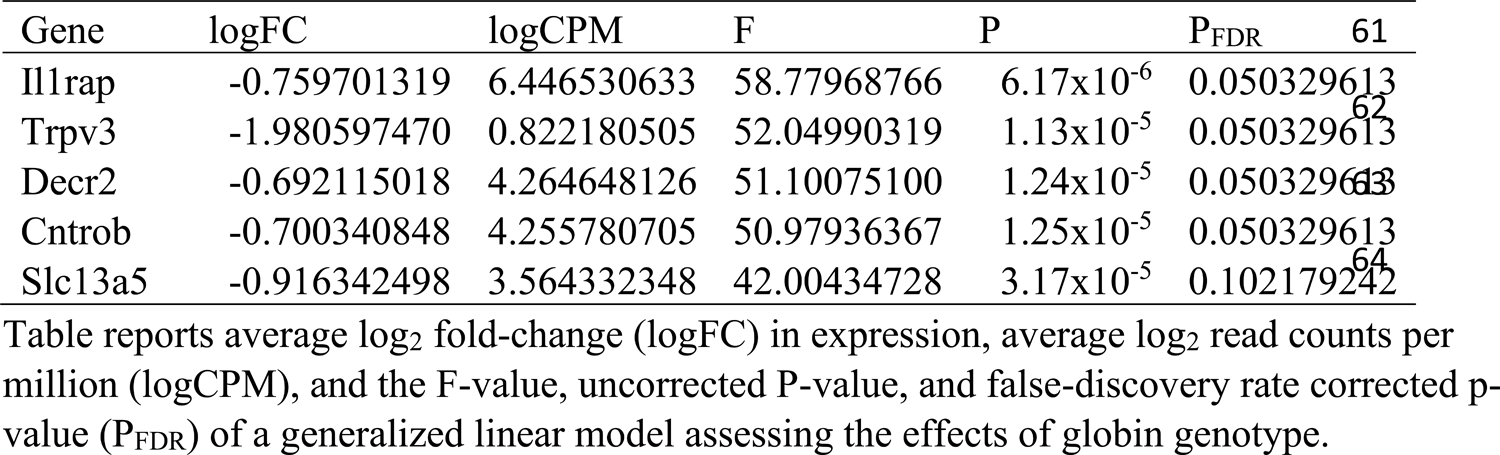
Genes that were differentially expressed in the medulla between α^HH^β^HH^ mice and α^LL^β^HL^ mice at P<0.10 after correction for the false discovery rate (P_FDR_).

Table S11. Comparison of modules from WGCNA between genotypes. Table reports uncorrected p-values, and false-discovery rate corrected p-values (PFDR).

